# *in silico* analysis and comparison of the metabolic capabilities of different organisms by reducing metabolic complexity

**DOI:** 10.1101/2025.08.08.669321

**Authors:** Evangelia Vayena, Meriç Ataman, Vassily Hatzimanikatis

**Affiliations:** Laboratory of Computational Systems Biotechnology, École Polytechnique Fédérale de Lausanne, EPFL, Lausanne, Switzerland

**Keywords:** microbiome modeling, comparative functional analysis, metabolic network reduction

## Abstract

Understanding how metabolic capabilities diverge across microbial species is fundamental for deciphering community function, ecological interactions, and for guiding synthetic microbiome design. Despite shared core pathways, microbial phenotypes can differ markedly due to evolutionary adaptations and metabolic specialization. Genome-scale metabolic models (GEMs) provide a systems-level framework to explore these differences; however, their complexity poses significant challenges to direct comparison. Here, we introduce NIS, a computational approach that uses the redGEM, lumpGEM, and redGEMX algorithms to systematically reduce GEMs to interpretable modules. NIS enables direct comparison of fueling pathways, biosynthetic routes, and environmental exchange processes across organisms, while preserving key metabolic information. We demonstrate the utility of NIS by analyzing *Escherichia coli* and *Saccharomyces cerevisiae*, revealing both conserved and divergent patterns in central metabolism, biomass biosynthesis, and substrate utilization. We further apply NIS to members of the core honeybee gut microbiome, uncovering distinct metabolic traits and complementarity that explain coexistence and interaction potential. Our framework offers a robust and scalable method to dissect microbial metabolic networks and supports the rational design and ecological understanding of microbial communities.

## Introduction

Microorganisms exhibit a remarkable diversity of phenotypes, even when exposed to identical environmental conditions. This phenotypic variability is rooted in the evolutionary divergence of metabolic capabilities and cellular requirements, factors that govern how organisms process nutrients and build cellular biomass Understanding the metabolic basis of these phenotypes is crucial for deciphering microbial behavior, niche adaptation and ecological interactions.

In microbial ecosystems, the architecture of an organism’s metabolic network influences how it interacts with others. Metabolic overlap can lead to competition for shared substrates^1^, while complementary pathways may foster cooperative interactions, including cross-feeding^2^. Therefore, understanding metabolic similarities and differences across species is essential to explaining coexistence, niche partitioning, and the emergent functions of microbial communities.

Most comparative studies rely on genome content and pathway inference based on the presence or absence of annotated genes^3–6^. However, such gene-centric approaches often simplify metabolism by reducing it to linear pathways, thereby overlooking the complex interdependencies between reactions and pathways within a complex network^6^. This limits their ability to explain phenotypic variation, particularly in organisms that are less characterized, where annotations are incomplete or biased by canonical reference pathways.

Genome-scale metabolic models (GEMs) address many of the limitations of gene-based methods by combining genomic, biochemical, and experimental data to create comprehensive networks of metabolic reactions. GEMs have been successfully developed for model organisms and pathogens^7–10^, human cells^11,12^, and more recently, for large collections of environmental^13,14^ and human-associated microbes^15–17^. GEMs enable the simulation-based exploration of metabolic capabilities, nutrient requirements, and potential ecological roles.

However, GEMs are inherently complex, comprising hundreds to thousands of metabolites and reactions. Comparing such models across organisms is both computationally and conceptually challenging. Existing strategies often rely on perturbation-based analyses, such as measuring the impact of deleting a shared reaction on metabolite production rates^18^ or determining the flux adjustments needed to restore steady-state after perturbing common reactions^6^. While these methods provide insights into functional similarities, they are inherently restricted to the shared biochemistry of the models and typically require solving numerous optimization problems, making them computationally intensive. Other approaches have been developed to systematically identify metabolic capabilities, such as enumerating minimal sets of metabolic conversions that a network can carry out between input and output metabolites to reveal network functions^19^, or to reduce the complexity of GEMs in microbial communities for scalable analysis^20^. However, these methods do not integrate fundamental physiological concepts with model reduction to facilitate meaningful comparisons across species. Therefore, there is a need for a more systematic and unbiased framework that can reduce complexity while preserving mechanistic and functional context, enabling meaningful comparisons across diverse organisms.

Inspired by classical microbial physiology, cellular metabolism can be modularly decomposed into three functional domains: (i) fueling pathways, which convert environmental substrates into precursor metabolites and energy; (ii) biosynthetic pathways, which synthesize the building blocks of cellular biomass; and (iii) exchange pathways, which govern nutrient uptake and byproduct secretion. This functional partitioning provides a structured and biochemically grounded framework for comparing metabolic capabilities across organisms. In particular, Neidhardt, Ingraham, and Schaechter^21^ proposed that biosynthetic pathways comprise reactions that transform twelve key precursor metabolites into the major macromolecular components of the cell. Conversely, fueling pathways include those reactions that utilize precursors, nitrogen and sulfur sources, energy, and redox equivalents from available environmental substrates to support biosynthesis. Classifying reactions in this manner enables meaningful, system-level comparisons while preserving the underlying biochemical context.

To address the challenges outlined above, we introduce NIS (Neidhardt–Ingraham– Schaechter), a computational workflow that leverages classical physiological definitions and integrates model reduction and subsystem comparison to extract and compare the functional modules of GEMs. Building on the algorithms redGEM^22^, lumpGEM^23^ and redGEMX^24^, NIS identifies core metabolic networks, reconstructs biosynthetic subnetworks, and explores environmental exchange capabilities. NIS provides a scalable and unbiased method for comparing organisms at multiple levels of metabolic organization.

We applied NIS to systematically compare *Escherichia coli* and *Saccharomyces cerevisiae*, revealing how compartmentalization, network connectivity, and energetic demands shape central metabolism and biosynthetic strategies. We then extended the analysis to the core honeybee gut microbiome, uncovering key metabolic traits, auxotrophies, and potential interaction patterns among community members. These applications exemplify how NIS can provide ecological and engineering insights into microbial metabolism.

## Materials and Methods

### Overall workflow

To systematically compare the metabolic capabilities of different organisms using GEMs, we developed NIS, a three-step workflow (Figure 1) that can be applied to any desired system. The workflow integrates the published algorithms redGEM^22^, lumpGEM^23^, and redGEMX^24^ to systematically reduce GEMs to core metabolic models and thus allow their comparison.

**Figure 1:**
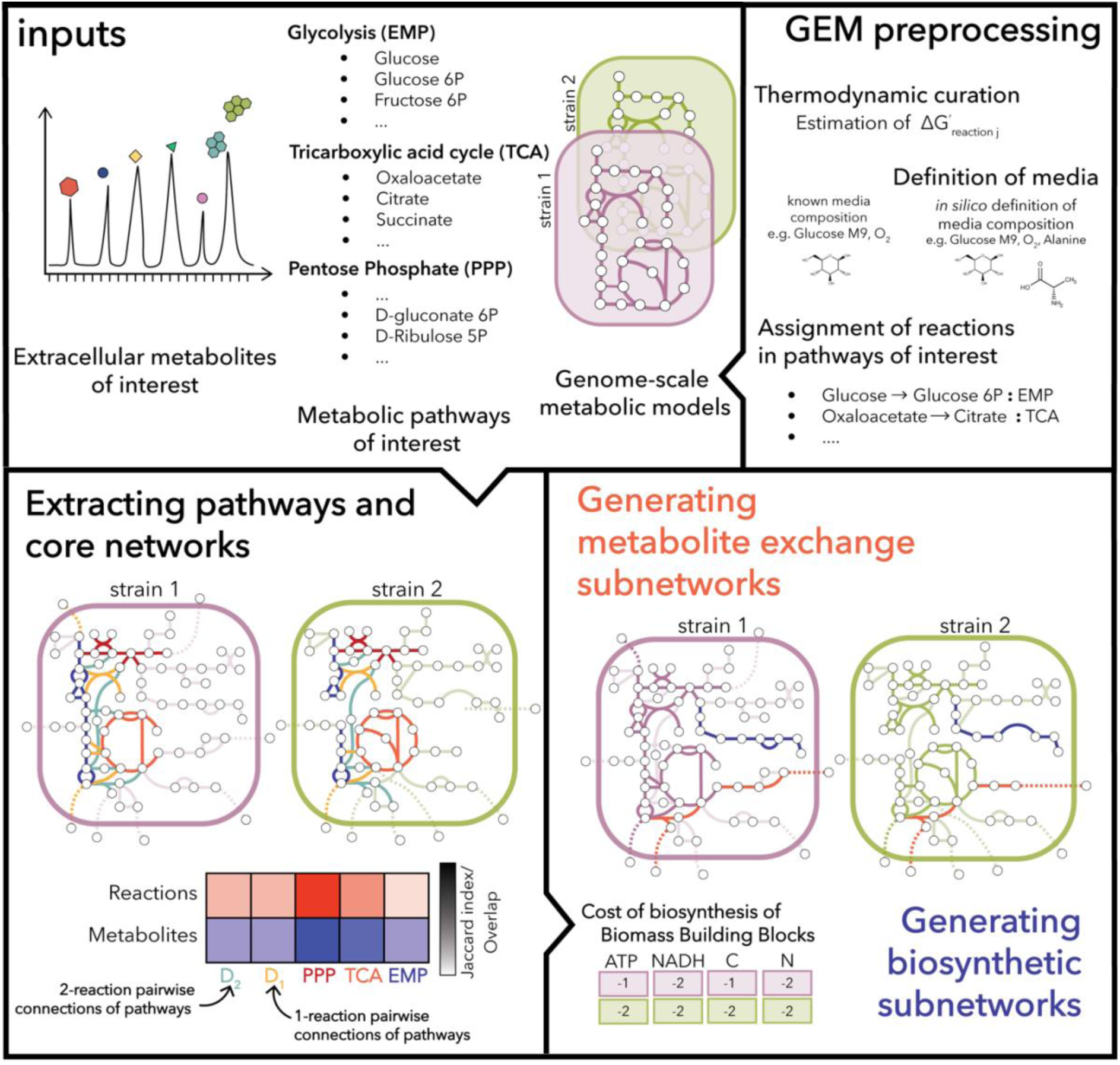
For the NIS workflow, metabolic subsystems and extracellular metabolites relevant to the study are selected. The GEMs are pre-processed (i.e., minimal media and assignment of reactions to subsystems) and thermodynamic constraints are added. A network expansion is then performed to connect the initial subsystems in a pairwise manner using reactions from the GEM to generate a core network for a selected degree of connectivity D. The minimal subnetworks required to connect the extracellular medium components to the core network and to produce the biomass building blocks are identified. At each step, the subnetworks, networks, and subsystems are compared.

We use as inputs: (i) the GEMs of the organisms we wish to compare; (ii) the metabolic subsystems that are of interest for the physiology under study; (iii) extracellular compounds that are relevant to the native environment of the organisms under study. To facilitate the comparison the GEMs should be pre-processed to have a harmonized reaction and metabolite nomenclature, with reactions consistently assigned to subsystems.

In the first step, we employ redGEM^22^. The algorithm first extracts the reaction and metabolite content of the subsystems of interest. Then, the initial network is expanded to include additional reactions from the GEM whose reactants and products belong to the initial subsystem. We can thus enrich the core network with reactions based on their metabolite content belonging to these subsystems but are not annotated as such in the reference pathways. Next, the algorithm performs a directed graph search to find the reactions from the GEM that connect in a pair-wise manner the initial metabolic subsystems for different degrees of connectivity D (e.g., D_1_, D_2_, D_3_ core networks). D represents the number of reactions that are used to connect pairs of metabolites from the different subsystems. We refer to the core network generated in the previous step as D_0_ core network. We then compare the metabolite and reaction content of the core networks (e.g., D_0_, D_1_ core networks). Comparisons may include or exclude intracellular compartmentalization, depending on the analysis. This choice depends on whether the physiological question requires capturing compartment-specific differences or focusing on the overall metabolic connectivity independent of localization.

In the second step, we employ lumpGEM^23^ on the D_0_ core networks and extract all alternative balanced subnetworks from the GEM to produce the biomass building blocks (BBBs) from precursors of the D_0_ core networks. We assess the similarity of the biosynthetic pathways for a given metabolite of the different organisms by comparing the union of all the alternative subnetworks generated from lumpGEM. The algorithm also lumps each one of the generated subnetworks into a single reaction. We use the stoichiometry of the lumped reactions to estimate the metabolic cost to produce each one of the BBBs. In the case of more than one lumped reaction, we average the stoichiometric coefficients to estimate the cost. For this step of the workflow, in the bee gut microbiome study, we used different minimal media for the different species since some of them have auxotrophies for certain biomass building blocks (mainly amino acids, nucleotides, and vitamins).

Finally, for the last step of the workflow, we employ redGEMX^24^. We generate all alternative balanced subnetworks from the GEM to connect selected extracellular metabolites to the core networks of the different organisms. We then compare the generated subnetworks. Alternatively, one can only focus on the nutrient uptakes and metabolic product secretions. This gives insight into the metabolic fate of the substrates without taking into account the different enzymes used to perform the biotransformation. In this step, a shared extracellular environment (union of all minimal media) was used for all bee gut microbiome organisms.

### GEM pre-processing

To compare the metabolism of *E. coli* and *S. cerevisiae*, we used the iML1515^7^ and Yeast8^8^ GEMs. As reactions in Yeast8 are often annotated to multiple subsystems, we used the procedure outlined below to systematically define subsystem content. Simulations were performed under M9 glucose minimal medium and aerobic conditions.

The core bee gut microbiome was defined based on existing studies^25–27^. The organisms included in this study are *Gilliamella apicola*, *Snodgrassella alvi*, *Bifidobacterium asteroides*, *Bombilactobacillus mellifer*, *Bombilactobacillus mellis*, *Lactobacillus apis* and *Lactobacillus* kullabergensis. Draft GEMs were acquired from the EMBL GEM collection (https://github.com/cdanielmachado/embl_gems), a collection of models built for all reference and representative bacterial genomes of NCBI RefSeq^28^ (release 84) using CarveMe^14^. Other core members (*L. helsingborgensis, L. kimbladii, L. melliventris*) were excluded due to lack of draft models. The *G. apicola*, *B. asteroides*, *L. apis* and *L. kullabergensis* GEMs were curated^29^ to capture anaerobic growth and/or transport of inorganics and vitamins and the *S. alvi* GEM to capture growth on minimal media. To constrain the extracellular environment of the cells, we performed an in silico minimal media analysis^24,30,31^ (Table S3). The CarveMe draft models do not contain information on the classification of reactions into metabolic subsystems. Thus, we used our in-house approach (see section “Metabolic subsystem definition” below) to define the reaction content of the subsystems of interest.

Thermodynamic curation of the GEMs, i.e., the Gibbs free energy of formation for the compounds in the GEMs and the corresponding error for the estimation, was performed using the Group Contribution Method. More specifically, MetaNetX^32^ (http://www.metanetx.org) was used to annotate the compounds of the GEM with identifiers from SEED^33^, KEGG^34–36^, CHEBI^37^, and HMDB^38^. Marvin (version 20.20, 2020, ChemAxon http://www.chemaxon.com) to transform the compound structures (canonical SMILES) into their major protonation states at pH 7 and to generate MDL Molfiles. We used the MDL Molfiles and the Group Contribution Method to estimate the standard Gibbs free energy of the formation of the compounds as well as the error of the estimation^39^. Furthermore, the thermodynamic properties of the different cellular compartments, including, pH, ionic strength, membrane potentials, and compartment concentration ranges have been integrated into the models as described before^40^.

### Metabolic subsystem definition

For the definition of the reaction content of the initial metabolic subsystems, i.e., glycolysis/gluconeogenesis, pentose phosphate pathway, tricarboxylic acid cycle, pyruvate metabolism, and oxidative phosphorylation the following procedure was followed. To define the reactions in the oxidative phosphorylation we extracted all reactions involving quinones and cytochromes. For the rest of the subsystems, we used the KEGG MODULES^34–36^ as a reference and we extracted the metabolite content of each module. More specifically, M00001, M00002, and M00003 were used to define the biotransformations of glycolysis/gluconeogenesis, M00004, M00005, M00008, M00166, and M00167 for the pentose phosphate pathway, M00009 for the tricarboxylic acid cycle and M00307, M00579, and M00172 for the pyruvate metabolism. We then used the ATLASx^41^ database to map the metabolite identify stereoisomers and map the metabolite IDs to different databases, e.g., the BiGG^9^database.

### Similarity assessment metrics

The Jaccard index^42^ was used as a metric to access the similarity of the core networks generated by redGEM and the pathways generated by redGEMX and lumpGEM. The Jaccard index, *J*, is a measure of the similarity between finite sample sets, A and B, and is defined as the size of the intersection divided by the size of the union of the sample sets:

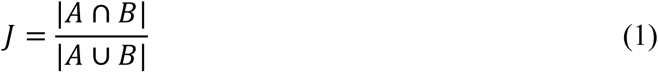

The index ranges from 0 to 1. The closer to 1, the more similar the two sets of data. The comparison was performed based on reaction or metabolite identifiers, i.e., sets A and B consisted of the reactions or metabolites of the network/subsystem under study of the two organisms under comparison.

## Results

### Comparative analysis of core metabolism in *E. coli* and *S. cerevisiae*

To evaluate the conservation and divergence of central carbon metabolism across phylogenetically distant microbes, we applied the NIS workflow to compare *Escherichia coli* (iML1515^7^ GEM) and *Saccharomyces cerevisiae* (Yeast8^8^ GEM) under aerobic glucose minimal conditions. Using a common subsystem definition for both organisms (see Materials and Methods), we extracted and analyzed five primary metabolic subsystems: glycolysis/gluconeogenesis, the pentose phosphate pathway, the tricarboxylic acid (TCA) cycle, pyruvate metabolism, and oxidative phosphorylation.

As a first step, we generated what we refer to as the D₀ core networks, which include only the reactions and metabolites assigned directly to these five subsystems. In the NIS framework, D₀ denotes the minimal core defined by subsystem membership, while D₁, D₂, …, Dᵢ progressively incorporate reactions needed to connect these subsystems through shortest paths of length i (in reaction steps), revealing degrees of connectivity^22^.

In the D₀ core networks, *E. coli* exhibited a more compact but highly connected network, with 147 reactions among 98 metabolites (Figure 2). In contrast, the *S. cerevisiae* D₀ network contained fewer reactions (112) but a larger set of metabolites (113) (Figure 2). This difference reflects both biological divergence and differences in network organization. *E. coli* possesses a more diverse array of quinone-based reactions due to its flexible electron transport system. In contrast, the eukaryotic compartments in *S. cerevisiae* (e.g., mitochondria and peroxisomes) expand the apparent metabolite set due to localization tags (Table S1).

**Figure 2:**
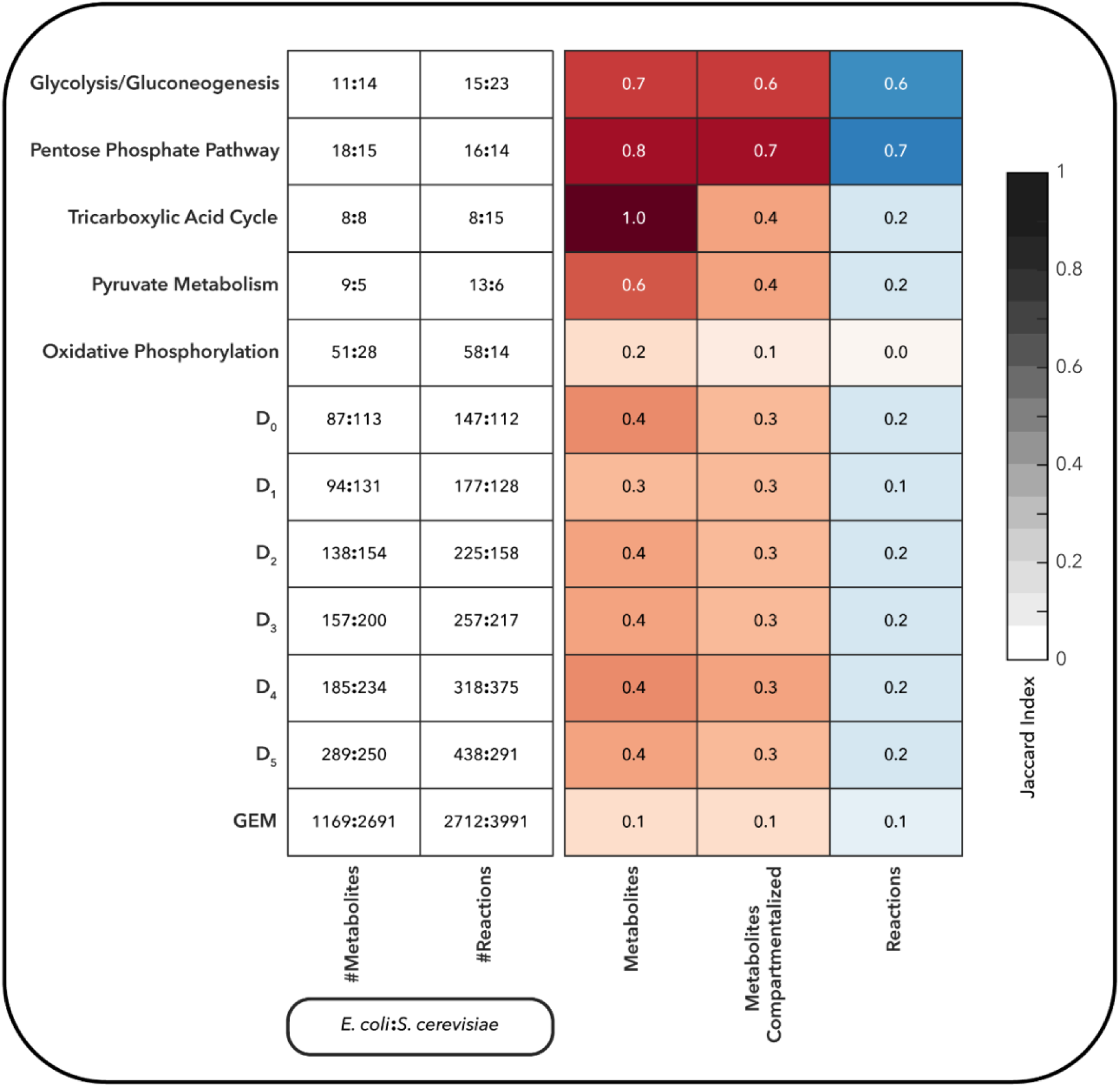
Comparison of metabolic subsystems, core networks, and GEMs of E. coli and S. cerevisiae, for the metabolite and reaction content.

We quantified similarity using the Jaccard index^42^ based on reaction and metabolite content. Glycolysis/gluconeogenesis and the pentose phosphate pathway showed the highest conservation, while oxidative phosphorylation and TCA-related reactions diverged substantially. Despite identical metabolite sets attributed to the TCA cycle (Jaccard index = 1 when compartments are ignored), the localization of these metabolites in mitochondria in *S. cerevisiae* versus the cytosol in *E. coli* leads to distinct reaction identifiers and network structures (e.g., the fumarate to malate biotransformation catalyzed by fumarase in noted as FUM in the iML1515 GEM whereas the same biotransformation is noted as FUMm in the Yeast8 GEM since it takes place in the mitochondria).

We further expanded the D₀ networks using redGEM to generate core networks for increasing degrees of connectivity D, from D₁ to D₅. Despite having more metabolites and a larger overall GEM, the *S. cerevisiae* core networks remained smaller with respect to reactions than those of *E. coli* across all levels, indicating a denser and more interconnected central metabolism in *E. coli* (Figure 2). Focusing on the D₁ networks, which capture the minimal number of reactions required to connect each subsystem pair, we observed that *E. coli* had more pairwise subsystem connections than *S. cerevisiae* (Figure S1). In *E. coli*, all subsystem pairs could be connected via a single reaction (minimum inherent distance = 1), whereas in *S. cerevisiae* at least two reactions were required to link the pentose phosphate pathway and the TCA cycle. The *E. coli* network offered multiple alternative reaction paths and metabolite bridges for this connection, including malate-pyruvate, oxoglutarate-pyruvate, and oxaloacetate-pyruvate (Figure S2), while in *S. cerevisiae* the connection relied on less redundant pathways involving oxaloacetate and pentose intermediates such as ribose-5-phosphate or xylulose-5-phosphate, mediated by pyruvate (Figure S3). This suggests that *E. coli* central metabolism is not only smaller but also more redundantly connected, an architecture that may confer greater metabolic plasticity.

### Biosynthetic strategies and energetic costs of biomass building blocks

To assess how different organisms allocate metabolic resources toward biomass synthesis, we applied the second step of the NIS workflow to reconstruct and compare the biosynthetic subnetworks for biomass building blocks (BBBs) in *E. coli* and *S. cerevisiae*. Using the lumpGEM algorithm on each organism’s D₀ core network, we identified minimal and elementally balanced subnetworks that support the production of their respective biomass precursors.

The biomass formulation in Yeast8 contains 67 BBBs, whereas the iML1515 model of *E. coli* includes 95. The number of subnetworks generated for *E. coli* was significantly larger (3,133 total, of which 821 were unique in terms of lumped reactions) compared to *S. cerevisiae* (274 total, 258 unique in terms of lumped reactions), reflecting the higher pathway redundancy and flexibility of the bacterial network. Among the BBBs in the two models, 41 compounds are shared, with 10 being inorganic. For the remaining 31 organic BBBs, we systematically compared the biosynthetic routes and estimated the associated material and energy costs.

Our analysis revealed that amino acid biosynthesis is the most conserved category between the two organisms, both in terms of network structure and associated costs (Figure 3A-B, Table S2). Carbohydrates and cofactors, in contrast, showed greater variation. BBBs with similar biosynthetic routes, such as phenylalanine and serine, exhibited comparable mass and energy costs. Interestingly, certain compounds with divergent biosynthetic networks (e.g., L-glutamine) showed similar or even identical (e.g., L-valine) costs between the two organisms, suggesting functional convergence despite mechanistic differences. In many cases, these differences can be attributed to compartmentalization effects in *S. cerevisiae*, where reactions are localized to mitochondria or other organelles.

**Figure 3:**
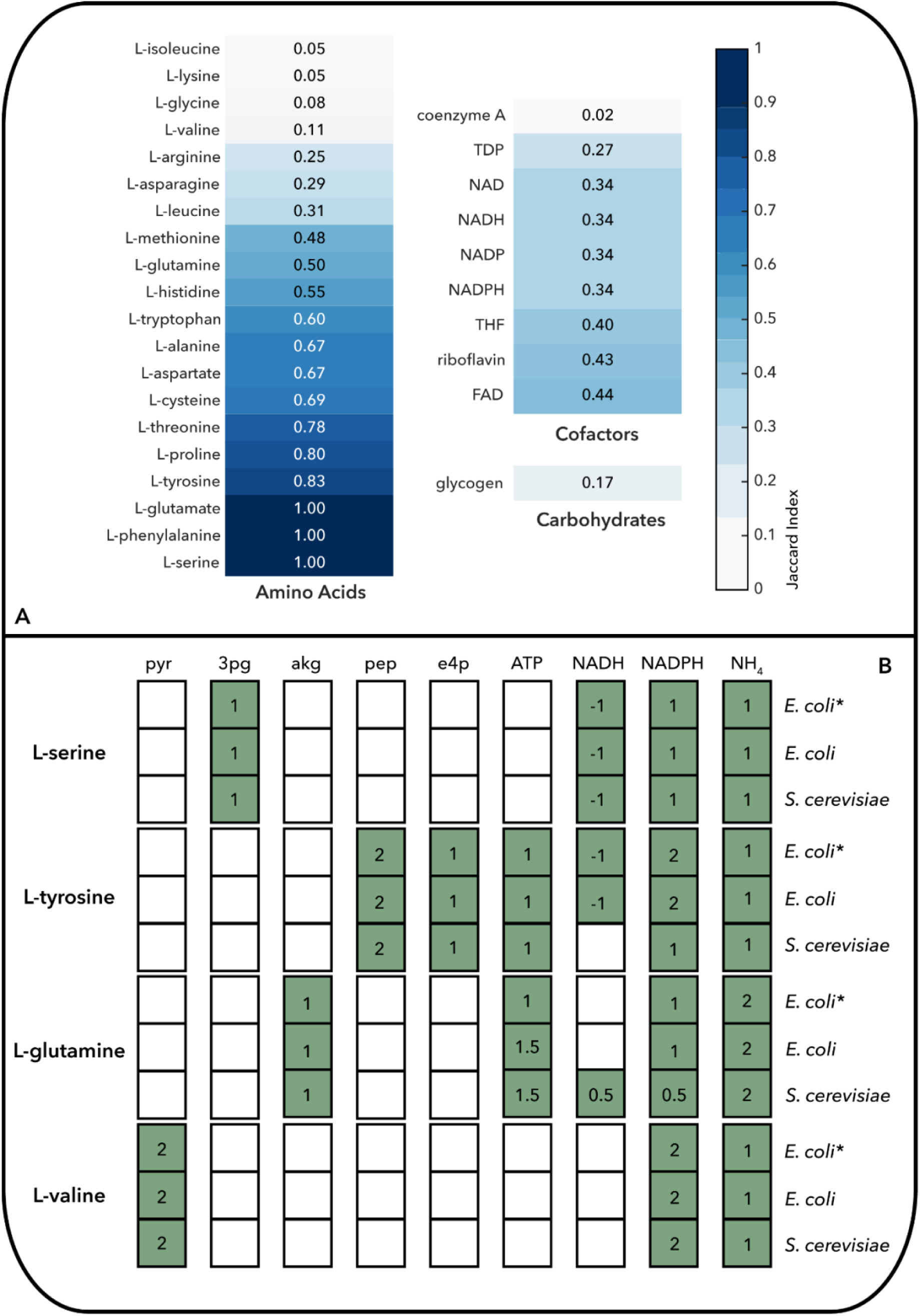
(A) Comparison of the subnetworks for the biosynthesis of different biomass building blocks in the E. coli and S. cerevisiae networks. (B) The cost of biosynthesis for selected amino acids in the E. coli and S. cerevisiae networks and comparison with previously reported values (*)^18^.

To further investigate how cost shapes biomass composition, we examined the correlation between biosynthetic cost and the abundance of amino acids in the biomass formulation (Figures S5-S6). In both organisms, amino acids derived from precursors such as phosphoenolpyruvate, erythrose 4-phosphate, or 3-phosphoglycerate (e.g., L-phenylalanine, L-tyrosine, L-tryptophan, and L-serine) tended to be less abundant (in mmol/gDW). Conversely, those synthesized from pyruvate or TCA cycle intermediates (e.g., L-valine, L-leucine, L-aspartate, and L-glutamate) were more prevalent. Amino acids whose biosynthesis requires more ATP, NADPH, and NADH units (e.g., L-methionine, L-arginine, and L-histidine) are generally less abundant, reinforcing the link between energetic investment and biomass allocation. This inverse relationship with biosynthetic cost is consistent with metabolic economy principles and evolutionary selection for efficiency. Notably, highly abundant amino acids such as L-alanine, L-aspartate, L-glutamate, and L-glycine in *S. cerevisiae* can be synthesized within one reaction step from the initial D₀ core network, underscoring the tight coupling between central metabolism and biomass demands. A similar pattern was observed for L-alanine in *E. coli*.

### Metabolite exchange pathways and substrate utilization strategies

To compare how different organisms interact with their environment through metabolic uptake and secretion, we applied the final step of the NIS workflow to analyze the exchange pathways in *E. coli* and *S. cerevisiae*. Using redGEMX, we reconstructed the minimal elementally balanced subnetworks required for each organism to catabolize a set of ten representative extracellular carbon compounds: glucose, acetate, lactate, pyruvate, succinate, fumarate, ethanol, malate, formate, and oxoglutarate^22^.

Although both organisms were able to uptake and catabolize all of the selected compounds in a M9 minimal media background and aerobic conditions, the minimal subnetworks through which each compound was metabolized differed substantially in structure (Figure 4A). This indicates divergence not only in transporter usage and reaction sequences, but also in how these compounds are functionally integrated into the core metabolism.

Despite differences in reaction content, we observed a high degree of convergence at the level of metabolic byproducts and co-substrate usage (Figure 4B). For six out of the ten compounds, lactate, oxoglutarate, succinate, ethanol, formate, and pyruvate, the identities of required co-substrates and resulting secreted byproducts were highly similar (Jaccard index ≥ 0.75). This suggests that, although the internal transformations differ, the functional metabolic fates of many compounds are conserved, possibly reflecting underlying physiological constraints.

Closer inspection of substrate-specific pathways highlighted key mechanistic differences (Figure 4C). For instance, glucose is fermented to lactate in *E. coli* via pyruvate, whereas *S. cerevisiae* converts glucose to ethanol. Acetate is catabolized to formate by *E. coli* using pyruvate formate lyase, a reaction absent in yeast. Instead, *S. cerevisiae* channels acetate through the glyoxylate shunt. Similarly, malate is transformed to pyruvate and formate in *E. coli,* but enters the TCA cycle in *S. cerevisiae* and is oxidized to oxaloacetate via malate dehydrogenase. These differences underscore how the presence or absence of key enzymes shapes environmental resource utilization, even under similar growth conditions.

**Figure 4:**
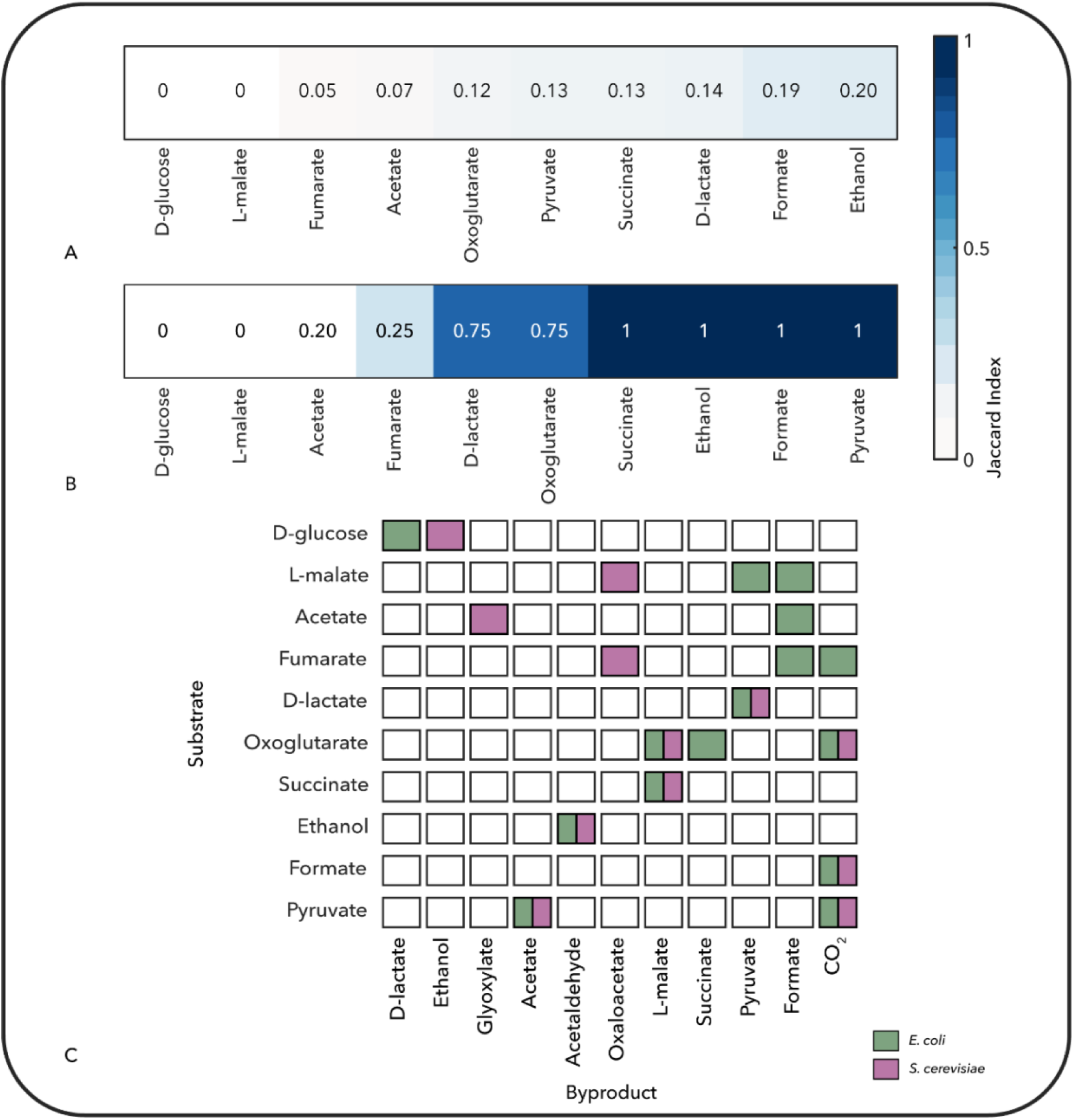
Comparison of (A) the subnetworks for the uptake and (B, C) the metabolic products of the catabolism of the ten carbon sources in the E. coli and S. cerevisiae networks.

**Figure 5:**
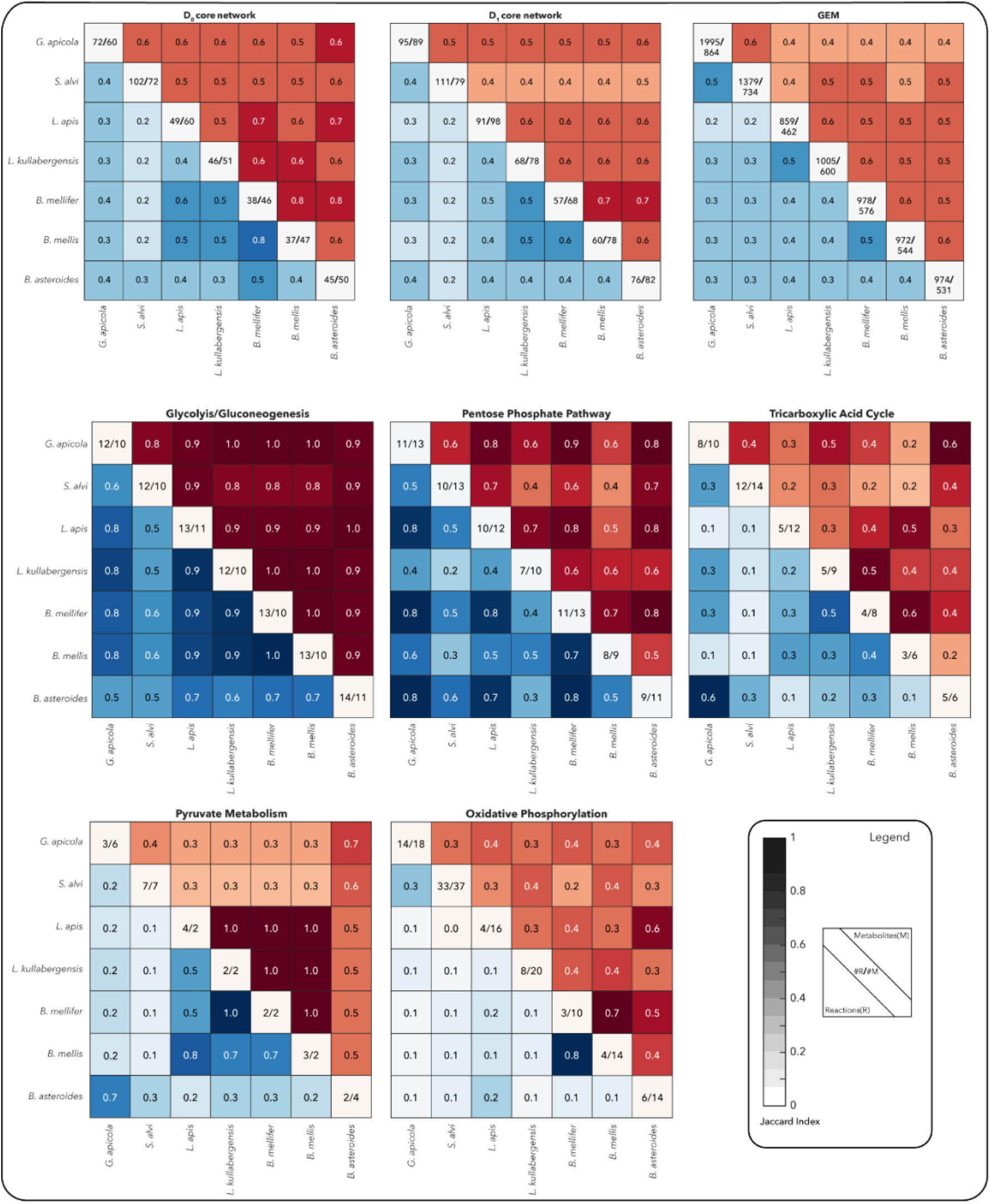
Pairwise comparison of metabolic subsystems, networks, and GEMs for the seven-member core honeybee gut microbiome, for the metabolite and reaction content.

### Metabolic diversity and niche differentiation in the core honeybee gut microbiome

To illustrate the ecological relevance and scalability of the NIS workflow, we applied it to the core honeybee gut microbiome. Extensive studies have been conducted around honeybees, due to their role as agricultural pollinators and their complex social behavior. Furthermore, over the past decade, the honeybee has been widely used as a model organism to study the gut microbiome^25^. The honeybee microbiome shows similarities with the human microbiome but, unlike humans, honeybees have a simpler and well-defined microbiome consisting of five core classes of organisms that are cultivable in the lab and genetically tractable^25^. These features make the honeybee an attractive system for mechanistic microbiome studies, enabling precise dissection of community assembly and host-microbe interactions.

We analyzed the metabolic capabilities of seven dominant species: *Gilliamella apicola* and the obligate aerobe *Snodgrassella alvi* both colonizing the ileum; *Bifidobacterium asteroids* that is mainly located in the rectum; and *Bombilactobacillus mellifer* and *Bombilactobacillus mellis* (previously classified as *Lactobacillus*^27^) as well as *Lactobacillus apis*, and *Lactobacillus kullabergensis* that colonize the ileum and the rectum^26,43^. Using GEMs from the CarveMe pipeline^14^, we systematically compared these organisms across three levels of metabolic organization: core metabolism, biomass biosynthesis, and environmental exchange.

### Core metabolism reveals divergent architectures despite phylogenetic closeness

We first compared the core metabolic networks (D₀) of the seven species. The glycolysis/gluconeogenesis and the pentose phosphate pathway were well conserved among most pairs, while TCA cycle components, pyruvate metabolism, and oxidative phosphorylation showed considerable divergence (**Error! Reference source not found.**). Notably, strains from the same phylotype, such as *B. mellifer* and *B. mellis* (firm-4), exhibited high core network similarity (Jaccard index = 0.8), while others like *L. apis* and *L. kullabergensis* (firm-5) were surprisingly dissimilar. This highlights that taxonomic relatedness based on 16S rRNA is not necessarily predictive of metabolic overlap.

Although *B. mellifer* and *B. mellis* shared a similar D₀ core, their D₁ and GEM-level networks diverged, indicating that peripheral pathway differences can impact full-network connectivity. These findings underscore the value of mechanistic, model-based comparisons for uncovering functional diversity that escapes detection through gene-based or taxonomic profiling alone.

Interestingly, the high similarity observed in glycolytic and energy metabolism pathways among the *Lactobacillus* and *Bombilactobacillus* strains suggests potential competition for shared substrates under natural conditions^44,45^. In contrast, high dissimilarity in central metabolic pathways among the remaining species pairs may reflect niche complementarity or cross-feeding potential, consistent with previous ecological interpretations of metabolic dissimilarity as a proxy for reduced competition and increased cooperation^45,46^.

### Biosynthetic pathways reflect auxotrophies and cross-feeding potential

We next compared the biosynthetic subnetworks for 40 organic biomass building blocks. In this analysis, all models shared the same biomass formulation, which was generated using the CarveMe workflow^14^. The most conserved biosynthetic patterns were observed in the amino acid category, particularly for the *S. alvi-G. apicola* pair, followed by *G. apicola-B. asteroides* and *S. alvi-B. asteroides* (Figure 6). Both *S. alvi* and *G. apicola* could synthesize all amino acids de novo, and their biosynthetic subnetworks showed high overlap (Jaccard index ≥ 0.7). These two species represent the only members of the community with fully complete amino acid biosynthesis capabilities. On the contrary, the nucleotide biosynthesis shows low similarity for the *S. alvi*-*G. apicola* pair, since *G. apicola* lacks enzymes for synthesizing the pyrimidines^47^ (in this study CMP is provided in the minimal media to simulate growth).

**Figure 6:**
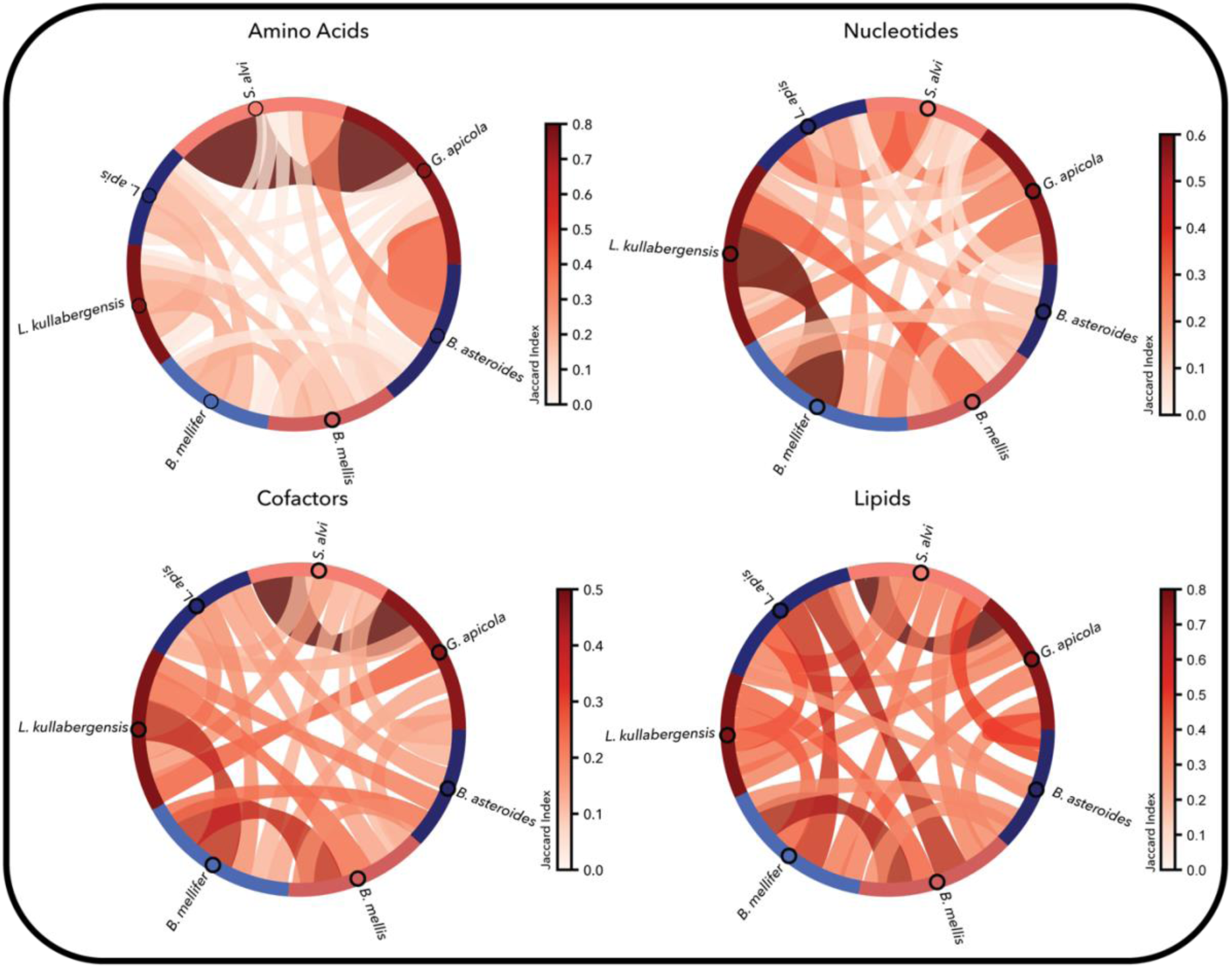
Comparison of the biosynthetic pathways for the biomass building blocks in the seven-member bee gut microbiome. We grouped the biomass building blocks in four categories and we present here the pairwise average similarity per category. The highest similarity is observed for the G. apicola-S. alvi pair and the amino acid biosynthesis.

The remaining species displayed distinct auxotrophy patterns. The firm-5 strains (*L. apis*, *L. kullabergensis*) were auxotrophic for the majority of amino acids, followed by the firm-4 strains (*B. mellifer*, *B. mellis*), which lacked several biosynthetic pathways as well. *B. asteroides* retained more capabilities but was still auxotrophic for L-cysteine and the aromatic amino acids (Table S3). These deficiencies suggest a reliance on nutrient acquisition from the environment or via cross-feeding within the community.

The diversity in biosynthetic network structure is also reflected in the estimated costs of amino acid production across species (Figure 7 and Table S4). In strains without auxotrophies, these costs were generally similar across organisms. However, in auxotrophic strains, the cost of biosynthesis became dependent on additional environmental metabolites beyond the canonical twelve precursors^14,21^. Although some biotransformations are still required to generate specific compounds (e.g., L-glutamate is a precursor to L-glutamine), the absence of full biosynthetic pathways leads to lower overall demands for energy and redox cofactors. This reduction in biosynthetic burden reflects not only a decrease in material requirements from core carbon and energy sources, but also highlights the impact of auxotrophy in shaping metabolic economy and potential interspecies dependencies.

**Figure 7:**
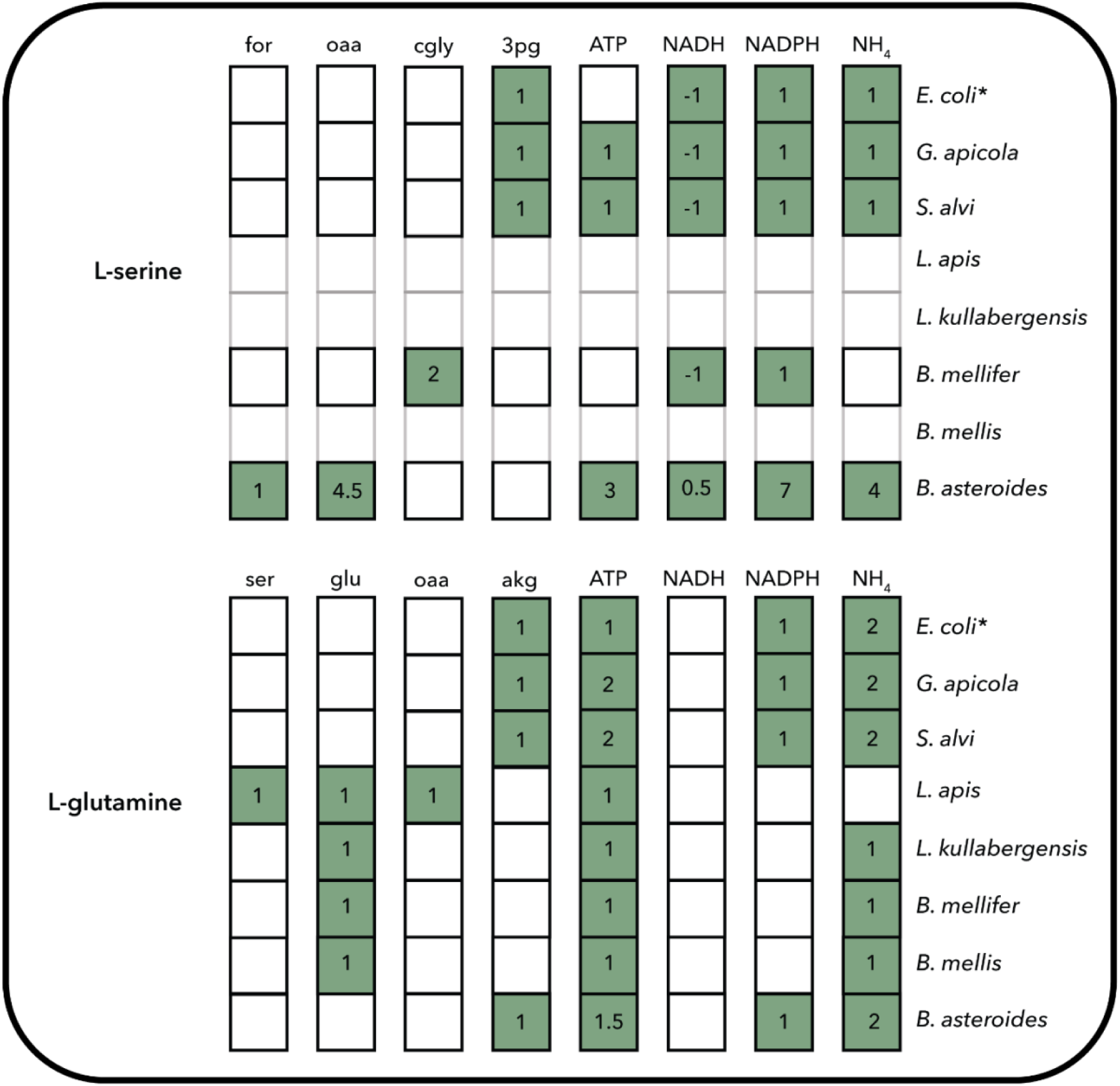
The cost of biosynthesis for selected amino acids in the core bee gut microbiome and comparison with previously reported values (*)^18^ for E. coli.

### Environmental exchange profiles differentiate substrate utilization strategies

Finally, we evaluated the ability of each strain to metabolize or secrete a curated pool of extracellular metabolites. We used the same pool of extracellular metabolites as in the *E. coli-S. cerevisiae* study above, and we also added citrate to this pool, since it has been suggested to be utilized by members of this community^15,46^. We found broad differences in exchange capabilities (Figure 8A). *S. alvi* and *G. apicola* had the most versatile profiles, capable of metabolizing nearly all compounds tested, followed by the *Lactobacillus* firm-5 members which are connected with 9 metabolites each. In contrast, *B. mellis* showed the most restricted exchange network and could secrete only succinate under the simulated environment. Even among close relatives, notable differences emerged, e.g., *B. mellifer* could metabolize TCA intermediates and secrete multiple compounds, whereas *B. mellis* could not.

**Figure 8:**
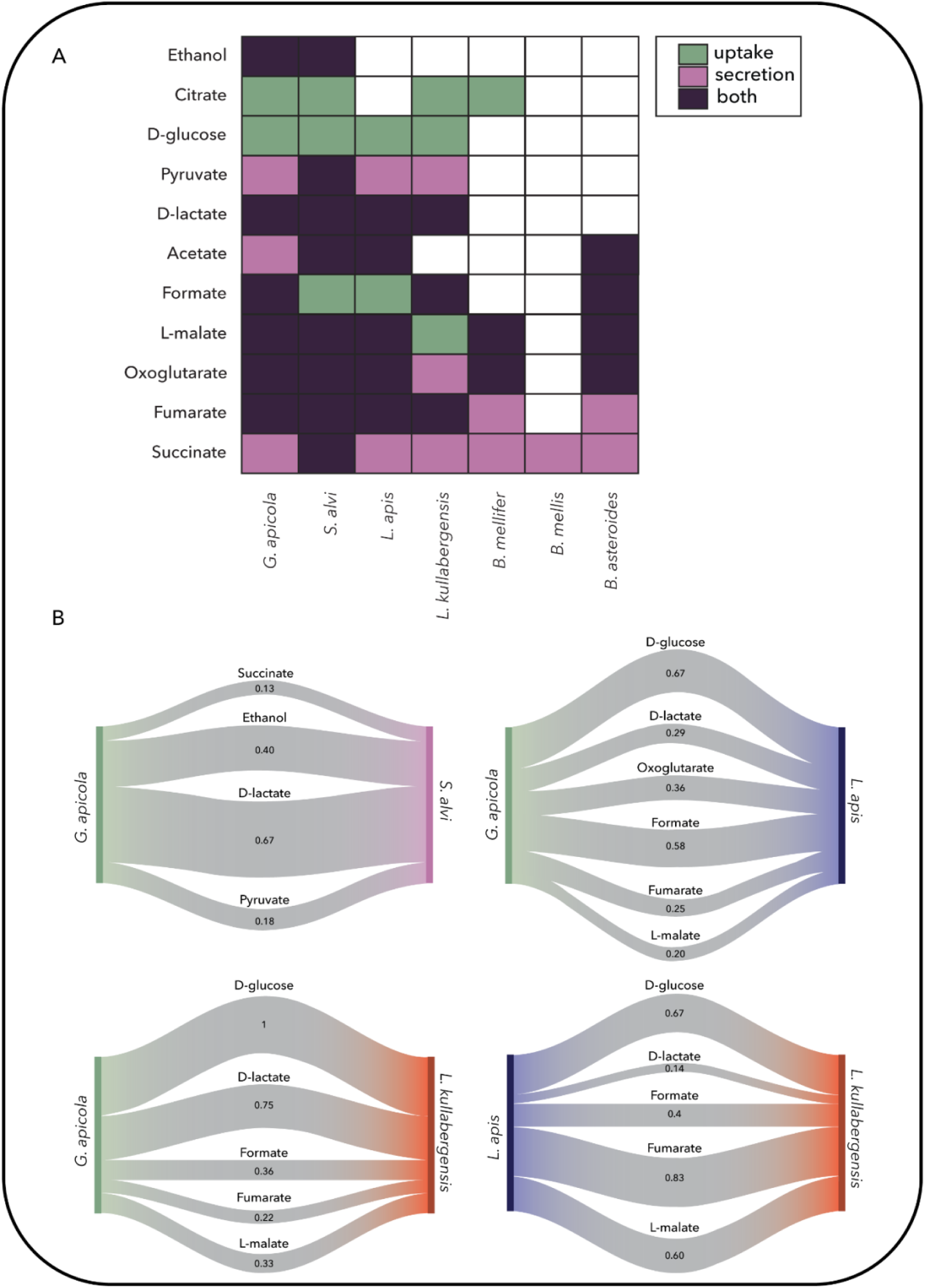
(A) The ability of the seven-member bee gut microbial community to uptake and/or produce different metabolites. (B) Comparison of the metabolic fate of different carbon sources for selected strain pairs. For each pair of organisms, each string corresponds to the compound it is labeled with, and the width of the string is proportional to the value of the Jaccard index that is also provided for each string.

Subnetwork comparisons (Figure 8B) revealed that similar metabolic outputs, such as lactate or malate secretion, can arise from distinct internal pathways, co-substrate requirements, and network structures. For instance, all core honeybee gut strains are capable of producing succinate, yet only *S. alvi* can further degrade it to malate via fumarate. Citrate is another key substrate that illustrates divergent processing. *S. alvi* utilizes it directly as a carbon source, producing oxoglutarate and CO₂, whereas *B. mellifer* requires co-metabolism with fructose and secretes malate, acetaldehyde, and CO₂. *L. apis* and *L. kullabergensis* share similar fumarate catabolism pathways. *L. apis* requires glucose co-uptake and secretes D-lactate and CO₂, while *L. kullabergensis* additionally secretes L-lactate. Similarly, *G. apicola* and *L. kullabergensis* exhibit comparable lactate utilization patterns, though their substrate preferences differ. *G. apicola* uses glucose or fructose, while *L. kullabergensis* is restricted to fructose. These context-dependent transformations underscore how niche specialization, co-substrate dependencies, and differential network connectivity shape potential metabolic cooperation within the community.

## Discussion

Microbial phenotypes are shaped not only by genomic content but also by the structure, connectivity, and utilization of metabolic networks under specific environmental conditions. In this study, we introduce and apply the NIS workflow, a computational framework designed to systematically compare microbial metabolism across multiple levels of biochemical organization. By reducing the complexity of genome-scale metabolic models and extracting functionally defined modules, including core metabolism, biosynthetic capacity, and environmental exchange, NIS enables mechanistically grounded comparisons between organisms with divergent evolutionary histories or ecological roles. This approach provides insights into how metabolic architecture underpins phenotypic variation and offers a scalable foundation for studying microbial interactions, community assembly, and metabolic specialization.

Our results demonstrate that *E. coli* and *S. cerevisiae*, representing prokaryotic and eukaryotic systems respectively, exhibit notable and expected differences in how central metabolism is organized, connected, and integrated into biomass formation and substrate use. The denser and more redundant architecture *of E. coli* central metabolism reflects its ecological strategy as a rapid responder and generalist, while the compartmentalized, less interconnected metabolism of *S. cerevisiae* reflects constraints and specializations linked to eukaryotic structure and fermentative capacity. Despite these differences, we observed convergence in biosynthetic costs, highlighting evolutionary pressures for metabolic efficiency.

Extending the analysis to the honeybee gut microbiome revealed how metabolic traits shape microbial coexistence. NIS uncovered both competitive and cooperative potentials based on biosynthetic overlap and substrate usage. In *S. alvi* and *G. apicola*, many amino acid biosynthesis pathways are conserved, whereas *Lactobacilli* and *Bombilactobacilli* show extensive auxotrophies. This contrast suggests that the latter species may depend on metabolic interdependence and cross-feeding within the community to acquire essential amino acids. We interpret the similarity in energy-generating pathways as competition, whereas the divergence in biosynthetic capabilities points to cooperation or neutral interactions. These insights expand our understanding of microbial association patterns and support the rational design of synthetic consortia.

NIS also opens avenues for translational applications. In medicine, characterizing metabolic dissimilarity may help identify commensal strains that can outcompete or complement pathogens. In agriculture and environmental microbiology, it can provide a framework for designing functionally robust microbial consortia. Beyond cross-species comparisons, NIS can also be applied to assess metabolic differences between GEMs of the same organism, capturing variations due to strain-level divergence, environmental adaptation, or modeling assumptions. The ability to compute biosynthetic costs and track substrate fate enables predictions about metabolic bottlenecks, dependencies, and resilience under perturbation. More broadly, NIS facilitates hypothesis generation at the intersection of systems biology and microbial ecology, where model-driven comparisons have long been constrained by scale and complexity.

Integrating NIS with experimental phenotyping, metagenomics, and multi-omics will allow more dynamic insights into how metabolism evolves within microbial communities. Future developments might include the integration of regulatory constraints, dynamic flux analysis, or community-level modeling to improve predictive power in complex environments. Nonetheless, by using metabolic principles and simplifying complex networks in a structured way, NIS offers a useful tool for both basic and applied microbiome research.

## Software

The simulations of this article were performed on Mac Pro 32 GB in MATLAB 2017a and IBM ILOG Cplex 12.7.1 as a solver.

## Acknowledgments

Funding for this work was provided by the Swiss National Science Foundation (SNSF): grant 200021_188623, NCCR Microbiomes, a National Centre of Competence in Research (grant number 180575), SystemsX.ch MicroScapeX grant 2013/158 and the École Polytechnique Fédérale de Lausanne.

## Author Contributions

EV, MA and VH conceptualized the study, developed the methodology and edited and reviewed the manuscript. EV and MA performed the data curation, developed the software, carried out the formal analysis, wrote the manuscript, developed the visualizations. VH managed and supervised the project. VH acquired the funding and the resources.

## Materials & Correspondence

Further information and requests for resources should be directed to and will be fulfilled by the corresponding author, Vassily Hatzimanikatis. The workflow and the models used in this study are available at the LCSB GitHub https://github.com/EPFL-LCSB/nis. The raw data is deposited on Zenodo^48^. The code for redGEM and lumpGEM is available at https://github.com/EPFL-LCSB/redgem and the code for redGEMX is available at https://github.com/EPFL-LCSB/redhuman.

