## Supplementary Material for "*in silico* analysis and comparison of the metabolic capabilities of different organisms by reducing metabolic complexity"

### Supplementary Figures

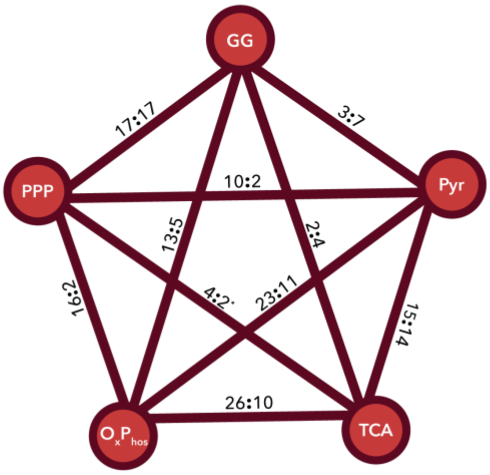


*Fig S 1: Number of alternative pairwise connections between the initial metabolic subsystems in the $D_1$ core network for E. coli : S. cerevisiae.*

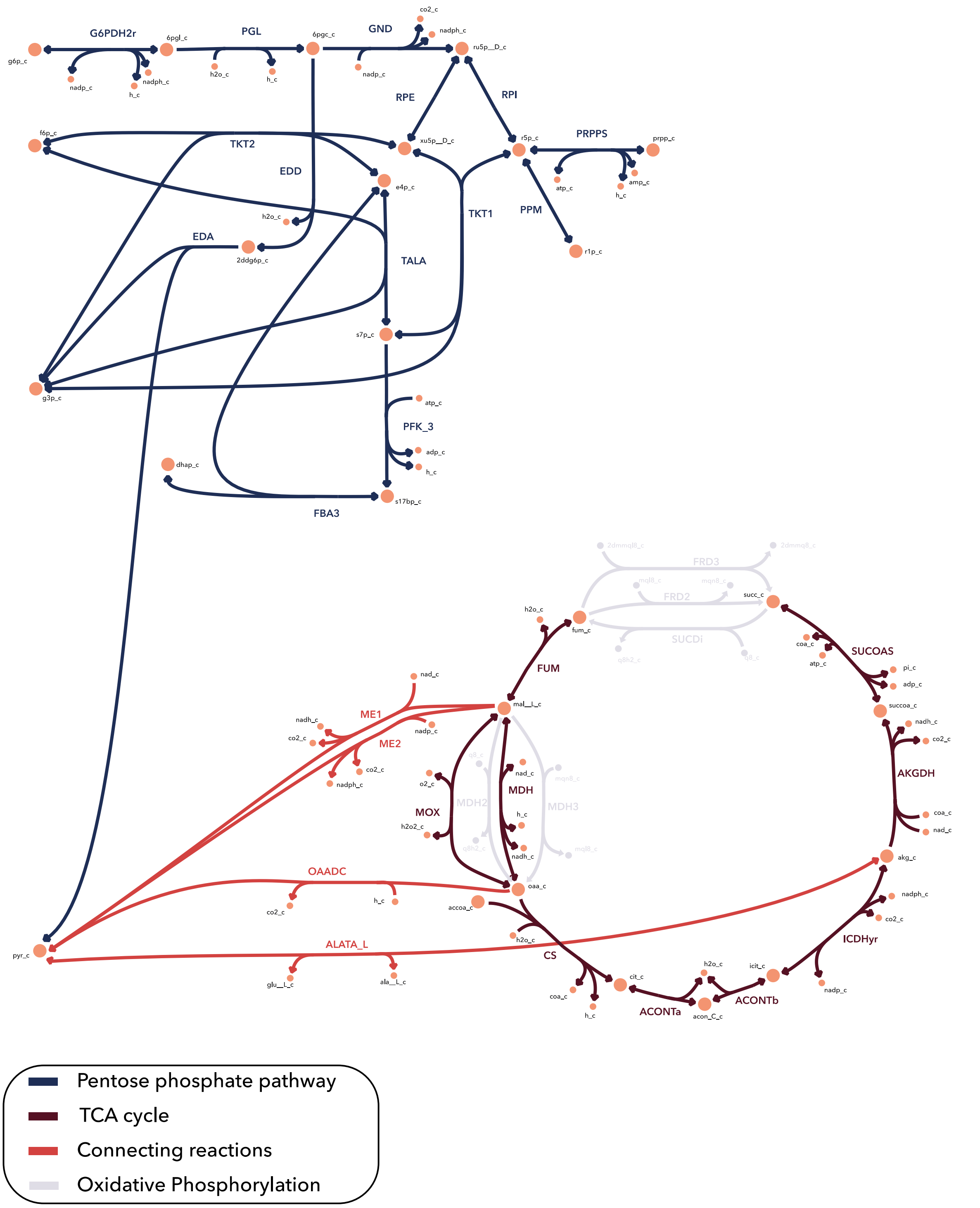


*Fig S 2: The pentose phosphate pathway and TCA cycle subsystems and the $D_1$ connections between them in the E. coli network. The plot is made with the Escher map web application[1].*

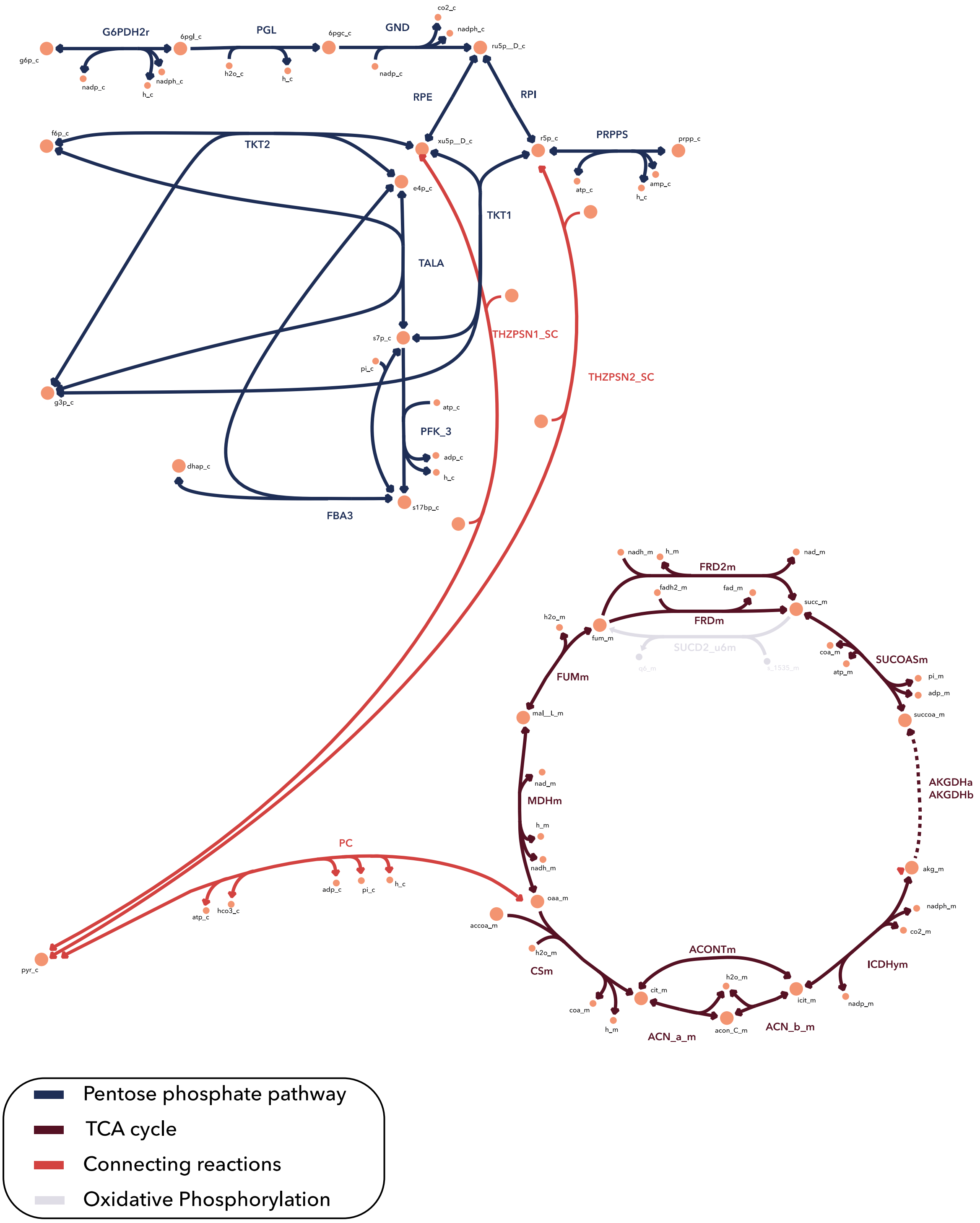


*Fig S 3: The pentose phosphate pathway and TCA cycle subsystems and the $D_1$ connections between them in the S. cerevisiae network. The plot is made with the Escher map web application[1].*

| | Minimal subnetwork size | | Alternatives | |
|---|---|---|---|---|
| | *E. coli* | *S. cerevisiae* | *E. coli* | *S. cerevisiae* |
| L-alanine | 2 | 2 | 1 | 2 |
| L-arginine | 11 | 12 | 1 | 4 |
| L-asparagine | 3 | 4 | 1 | 4 |
| L-aspartate | 2 | 2 | 1 | 2 |
| L-glutamate | 1 | 1 | 1 | 1 |
| L-glycine | 8 | 4 | 1 | 2 |
| L-isoleucine | 12 | 10 | 1 | 1 |
| L-leucine | 10 | 11 | 1 | 1 |
| L-lysine | 11 | 10 | 1 | 1 |
| L-phenylalanine | 11 | 11 | 1 | 1 |
| L-proline | 4 | 5 | 1 | 1 |
| L-serine | 4 | 4 | 1 | 1 |
| L-threonine | 7 | 7 | 1 | 4 |
| L-tryptophan | 14 | 17 | 1 | 2 |
| L-tyrosine | 11 | 11 | 1 | 1 |
| L-valine | 5 | 5 | 1 | 1 |
| glycogen | 3 | 4 | 1 | 1 |
| L-glutamine | 2 | 2 | 2 | 2 |
| L-cysteine | 13 | 12 | 3 | 1 |
| L-methionine | 23 | 19 | 3 | 4 |
| L-histidine | 18 | 28 | 6 | 4 |
| TDP | 43 | 41 | 6 | 61 |
| NAD | 28 | 44 | 12 | 4 |
| NADH | 28 | 44 | 12 | 4 |
| NADP(+) | 28 | 44 | 12 | 4 |
| NADPH | 28 | 44 | 12 | 4 |
| FAD | 36 | 33 | 24 | 16 |
| THF | 43 | 47 | 24 | 8 |
| riboflavin | 32 | 30 | 24 | 8 |
| coenzyme A | 45 | 6 | 36 | 1 |

*Fig S 4: The minimal subnetwork size for the biosynthesis of the common biomass building blocks in the E. coli and the S. cerevisiae networks and the number of alternative subnetworks of this size.*

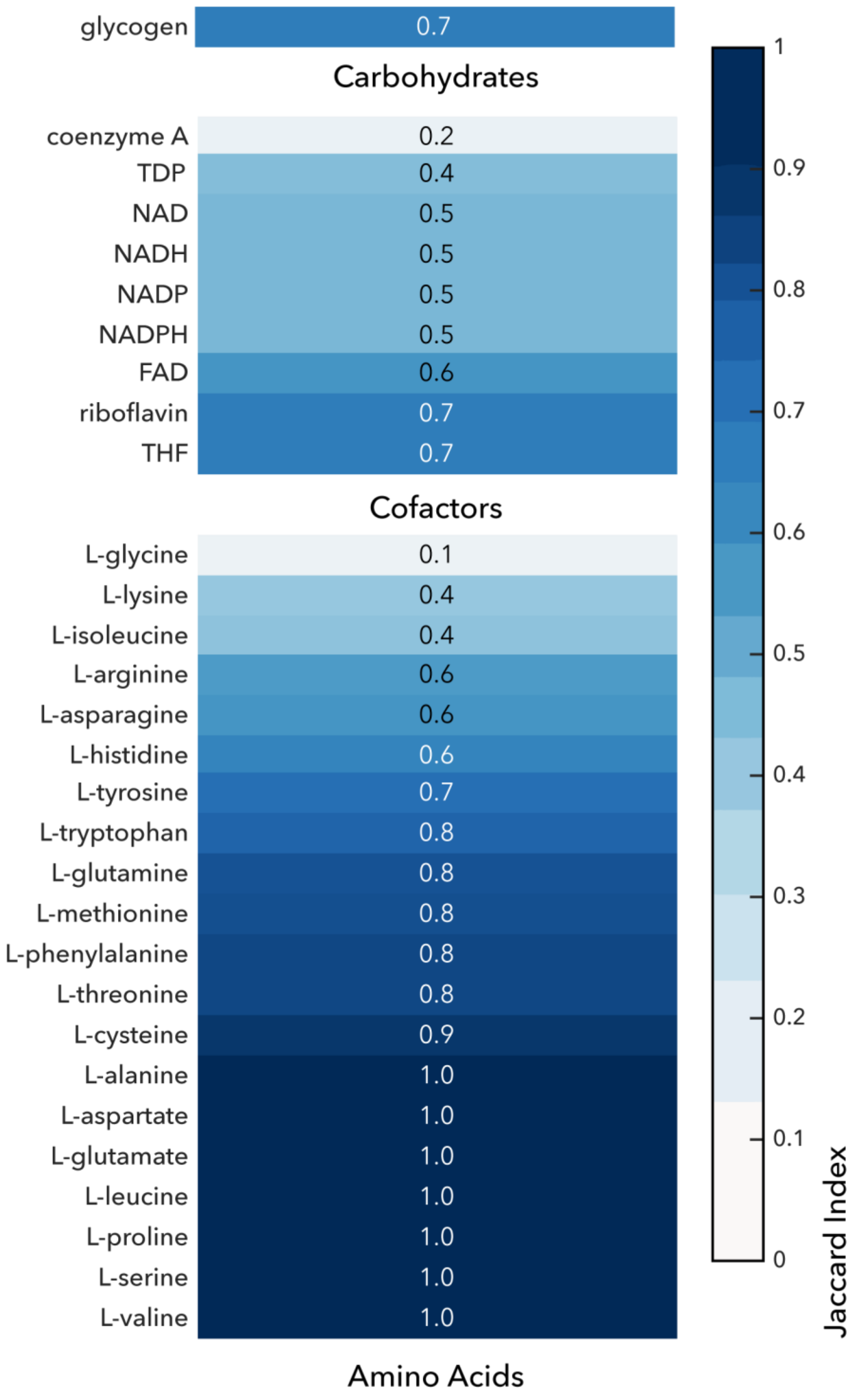


*Fig S 5: Similarity of the cost of biosynthesis of the common biomass building blocks in the E. coli and the S. cerevisiae networks.*

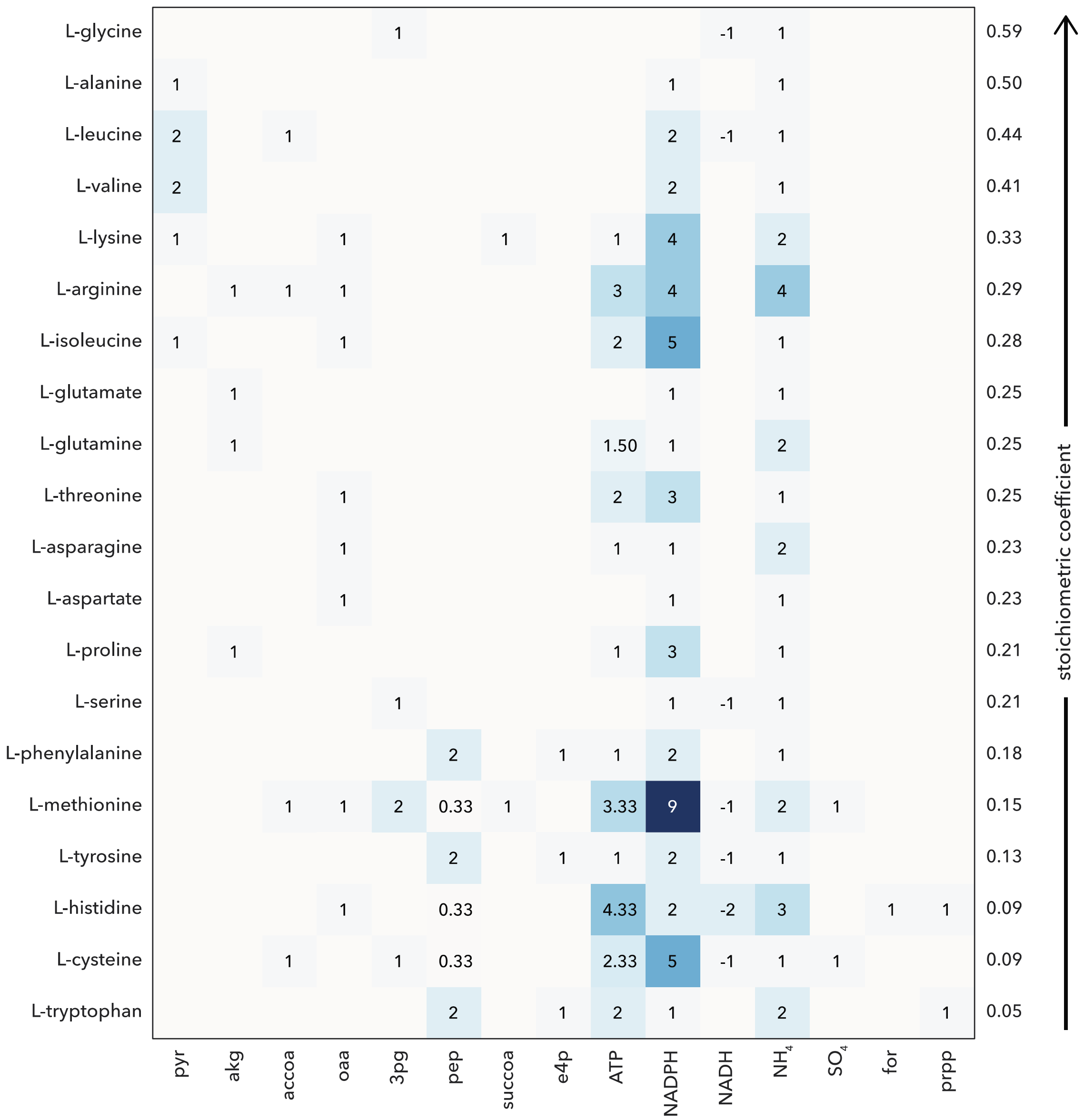


*Fig S 6: Cost of biosynthesis of the amino acids for E. coli. The amino acids are sorted based on their relative abundance in the biomass.*

| | pyr | akg | accoa | oaa | 3pg | pep | e4p | ATP | NADPH | NADH | $NH_4$ | $SO_4$ | icit | prpp | stoichiometric coefficient |
|---|---|---|---|---|---|---|---|---|---|---|---|---|---|---|---|
| L-alanine | 1 | | | | | | | | 1 | | 1 | | | | 0.53 |
| L-glutamate | | 1 | | | | | | | 1 | | 1 | | | | 0.35 |
| L-aspartate | | | | 1 | | | | | 1 | | 1 | | | | 0.34 |
| L-leucine | 2 | | 1 | | | | | | 2 | -1 | 1 | | | | 0.34 |
| L-glycine | | | | | | | | | 1 | | 1 | | 1 | | 0.33 |
| L-lysine | | 1 | 1 | | | | | 1 | 4 | -2 | 2 | | | | 0.33 |
| L-valine | 2 | | | | | | | | 2 | | 1 | | | | 0.30 |
| L-isoleucine | 1 | 1 | | | | | | | 2 | | 1 | | | | 0.22 |
| L-threonine | | | | 1 | | | | 2 | 2.50 | 0.50 | 1 | | | | 0.22 |
| L-serine | | | | | 1 | | | | 1 | -1 | 1 | | | | 0.21 |
| L-proline | | 1 | | | | | | 1 | 3 | | 1 | | | | 0.19 |
| L-arginine | | 1 | | 1 | | | | 6.50 | 2.50 | 1.50 | 4 | | | | 0.18 |
| L-phenylalanine | | | | | | 2 | 1 | 1 | 2 | | 1 | | | | 0.15 |
| L-glutamine | | 1 | | | | | | 1.50 | 0.50 | 0.50 | 2 | | | | 0.12 |
| L-tyrosine | | | | | | 2 | 1 | 1 | 1 | | 1 | | | | 0.12 |
| L-asparagine | | | | 1 | | | | 2.50 | 0.50 | 0.50 | 2 | | | | 0.12 |
| L-histidine | | | | 1 | 1 | | | 5.50 | 1.50 | -1.50 | 4 | | | 2 | 0.08 |
| L-methionine | | | 1 | 1 | 1 | | | 3 | 8.50 | -0.50 | 2 | 1 | | | 0.06 |
| L-tryptophan | | | | | 1 | 2 | 1 | 2.50 | 1.50 | -0.50 | 2 | | | 1 | 0.03 |
| L-cysteine | | | 1 | | 1 | | | 2 | 5 | -1 | 1 | 1 | | | 0.01 |

*Fig S 7: Cost of biosynthesis of the amino acids for S. cerevisiae. The amino acids are sorted based on their relative abundance in the biomass.*

| | Minimal subnetwork size | | Alternatives | |
|---|---|---|---|---|
| | *E. coli* | *S. cerevisiae* | *E. coli* | *S. cerevisiae* |
| Succinate | 9 | 12 | 1 | 1 |
| Ethanol | 14 | 14 | 1 | 1 |
| D-lactate | 13 | 15 | 1 | 1 |
| Formate | 13 | 16 | 1 | 1 |
| L-malate | 12 | 13 | 1 | 1 |
| Acetate | 14 | 21 | 2 | 1 |
| Oxoglutarate | 13 | 17 | 2 | 1 |
| Pyruvate | 13 | 16 | 2 | 1 |
| Fumarate | 13 | 11 | 3 | 1 |
| D-glucose | 9 | 9 | 6 | 1 |

*Fig S 8: The minimal subnetwork size for the uptake of ten selected compounds in the E. coli and the S. cerevisiae networks and the number of alternative subnetworks of this size.*

| | Minimal subnetwork size | | | | | | | Alternatives | | | | | | |
|---|---|---|---|---|---|---|---|---|---|---|---|---|---|---|
| | *G. apicola* | *S. alvi* | *L. apis* | *L. kullabergensis* | *B. mellifer* | *B. mellis* | *B. asteroides* | *G. apicola* | *S. alvi* | *L. apis* | *L. kullabergensis* | *B. mellifer* | *B. mellis* | *B. asteroides* |
| L-Alanine | 3 | 3 | 2 | 2 | 2 | 2 | 2 | 1 | 1 | 2 | 2 | 1 | 1 | 7 |
| L-Arginine | 14 | 12 | | | | 8 | 14 | 1 | 1 | | | | 1 | 8 |
| L-Aspartate | 3 | 3 | 3 | 3 | | | 2 | 1 | 1 | 1 | 2 | | | 2 |
| L-Glutamine | 2 | 2 | 4 | 1 | 1 | 1 | 2 | 1 | 1 | 2 | 2 | 1 | 1 | 4 |
| L-Glutamate | 2 | 2 | | | | | 1 | 1 | 1 | | | | | 2 |
| Glycine | 9 | 9 | 1 | 1 | 1 | | 8 | 1 | 2 | 2 | 2 | 1 | | 4 |
| L-Isoleucine | 13 | 12 | 7 | | 19 | | 12 | 1 | 2 | 4 | | 8 | | 4 |
| L-Lysine | 12 | 12 | 16 | 15 | 13 | 13 | 11 | 1 | 2 | 1 | 4 | 4 | 2 | 2 |
| L-Phenylalanine | 12 | 12 | | | | 10 | 7 | 1 | 1 | | | | 2 | 4 |
| L-Proline | 6 | 6 | 4 | 1 | | 10 | 5 | 1 | 1 | 2 | 2 | | 4 | 24 |
| L-Serine | 5 | 5 | | | 6 | | 12 | 1 | 1 | | | 1 | | 4 |
| L-Threonine | 8 | 8 | | | 13 | | 7 | 1 | 2 | | | 2 | | 4 |
| L-Tryptophan | 20 | 17 | | | | 13 | 24 | 1 | 1 | | | | 1 | 8 |
| L-Tyrosine | 12 | 12 | | | | 10 | 7 | 1 | 1 | | | | 2 | 4 |
| Pyridoxal 5'-phosphate | 2 | 8 | 7 | 3 | 4 | 2 | 2 | 1 | 1 | 2 | 2 | 1 | 1 | 6 |
| Thiamine diphosphate | 7 | 6 | 9 | 7 | 7 | 7 | 7 | 1 | 1 | 66 | 2 | 1 | 2 | 2 |
| L-Asparagine | 6 | 4 | 6 | 6 | 3 | 3 | 6 | 2 | 1 | 2 | 2 | 1 | 1 | 4 |
| L-Cysteine | 15 | 11 | 1 | 1 | 1 | 1 | 1 | 2 | 2 | 2 | 2 | 1 | 1 | 2 |
| L-Leucine | 11 | 11 | | | | | 10 | 2 | 1 | | | | | 8 |
| L-Valine | 6 | 6 | | | | | 5 | 2 | 1 | | | | | 10 |
| CTP | 13 | 12 | 16 | 8 | 9 | 22 | 3 | 3 | 1 | 4 | 2 | 1 | 2 | 2 |
| DTTP | 20 | 21 | 22 | 2 | 18 | 3 | 12 | 3 | 1 | 4 | 2 | 3 | 1 | 12 |
| UTP | 11 | 11 | 15 | 7 | 8 | 21 | 4 | 3 | 1 | 2 | 2 | 1 | 2 | 2 |
| L-Histidine | 21 | 42 | | | | | 21 | 6 | 4 | | | | | 4 |
| GTP | 29 | 31 | 25 | 9 | 9 | 6 | 9 | 6 | 4 | 2 | 2 | 1 | 1 | 4 |
| FAD | 40 | 35 | 24 | 24 | 13 | 10 | 5 | 6 | 4 | 1 | 4 | 1 | 1 | 2 |
| NAD | 32 | 37 | 28 | 15 | 12 | 9 | 4 | 6 | 8 | 2 | 2 | 1 | 1 | 2 |
| NADP | 32 | 37 | 28 | 15 | 12 | 9 | 4 | 6 | 8 | 2 | 2 | 1 | 1 | 2 |
| Riboflavin | 36 | 31 | | 19 | | | | 6 | 4 | | 8 | | | |
| DATP | 30 | 35 | 3 | 16 | 15 | 10 | 4 | 9 | 4 | 22 | 2 | 2 | 1 | 6 |
| DCTP | 15 | 14 | 20 | 13 | 13 | 24 | 5 | 9 | 1 | 4 | 4 | 1 | 2 | 6 |
| L-Methionine | 21 | 18 | 1 | | | | 14 | 12 | 16 | 1 | | | | 72 |
| 10-Formyltetrahydrofolate | 48 | 48 | 16 | 22 | 8 | 5 | 3 | 12 | 4 | 1 | 16 | 1 | 1 | 12 |
| 5,10-Methylenetetrahydrofolate | 50 | 50 | | 6 | 7 | 3 | 5 | 12 | 4 | | 6 | 1 | 1 | 12 |
| 5,6,7,8-Tetrahydrofolate | 48 | 48 | 2 | 5 | 4 | | 2 | 12 | 4 | 2 | 8 | 1 | | 12 |
| DGTP | 31 | 33 | 28 | 14 | 11 | 8 | 11 | 20 | 4 | 2 | 4 | 1 | 1 | 8 |
| uaagmda | 50 | 47 | 51 | 40 | 51 | 45 | 62 | 24 | 8 | 36 | 48 | 16 | 8 | 8 |
| S-Adenosyl-L-methionine | 40 | 43 | 24 | 15 | 12 | 9 | 18 | 72 | 64 | 1 | 2 | 1 | 1 | 72 |
| Menaquinol 8 | 56 | 48 | 35 | 34 | 91 | 37 | 38 | 216 | 4 | 6 | 1 | 3 | 8 | 48 |
| Coenzyme A | 55 | 55 | 31 | 25 | 34 | 40 | | 432 | 16 | 1 | 4 | 2 | 2 | |

*Fig S 9: The minimal subnetwork size for the biosynthesis of the biomass building blocks in the seven-member bee gut microbiome and the number of alternative subnetworks.*

**Minimal subnetwork size**

| | G. apicola | S. alvi | L. apis | L. kullabergensis | B. mellifer | B. mellis | B. asteroides |
|---|---|---|---|---|---|---|---|
| Pyruvate | | 10 | | | | | |
| Citrate | 8 | 12 | | 37 | 23 | | |
| Succinate | | 7 | | | | | |
| Ethanol | 14 | 11 | | | | | |
| D-glucose | 9 | 7 | 7 | 9 | | | |
| D-lactate | 13 | 9 | 17 | 11 | | | |
| Oxoglutarate | 16 | 10 | 24 | | 18 | | 17 |
| Fumarate | 14 | 10 | 16 | 14 | | | |
| Acetate | | 13 | 60 | | | | 19 |
| L-malate | 12 | 9 | 23 | 19 | 25 | | 20 |
| Formate | 22 | 11 | 21 | 22 | | | 25 |

**Alternatives**

| | G. apicola | S. alvi | L. apis | L. kullabergensis | B. mellifer | B. mellis | B. asteroides |
|---|---|---|---|---|---|---|---|
| Pyruvate | | 1 | | | | | |
| Citrate | 1 | 1 | | 7 | 1 | | |
| Succinate | | 2 | | | | | |
| Ethanol | 2 | 4 | | | | | |
| D-glucose | 2 | 1 | 1 | 10 | | | |
| D-lactate | 2 | 2 | 4 | 1 | | | |
| Oxoglutarate | 2 | 3 | 1 | | 2 | | 6 |
| Fumarate | 2 | 1 | 10 | 2 | | | |
| Acetate | | 6 | 12 | | | | 1 |
| L-malate | 6 | 2 | 2 | 2 | 8 | | 2 |
| Formate | 22 | 4 | 1 | 4 | | | 2 |

**Minimal subnetwork size**

| | G. apicola | S. alvi | L. apis | L. kullabergensis | B. mellifer | B. mellis | B. asteroides |
|---|---|---|---|---|---|---|---|
| Pyruvate | 15 | 9 | 20 | 18 | | | |
| Citrate | | | | | | | |
| Succinate | 14 | 10 | 24 | 19 | 18 | 42 | 17 |
| Ethanol | 10 | 14 | | | | | |
| D-glucose | | | | | | | |
| D-lactate | 9 | 7 | 7 | 8 | | | |
| Oxoglutarate | 13 | 11 | 23 | 19 | 20 | | 17 |
| Fumarate | 14 | 10 | 16 | 14 | 31 | | 27 |
| Acetate | 13 | 10 | 29 | | | | 17 |
| L-malate | 12 | 7 | 24 | | 23 | | 27 |
| Formate | 16 | | | 25 | | | 21 |

**Alternatives**

| | G. apicola | S. alvi | L. apis | L. kullabergensis | B. mellifer | B. mellis | B. asteroides |
|---|---|---|---|---|---|---|---|
| Pyruvate | 1 | 3 | 10 | 4 | | | |
| Citrate | | | | | | | |
| Succinate | 2 | 2 | 1 | 12 | 2 | 27 | 3 |
| Ethanol | 2 | 3 | | | | | |
| D-glucose | | | | | | | |
| D-lactate | 2 | 1 | 1 | 1 | | | |
| Oxoglutarate | 2 | 1 | 2 | 2 | 2 | | 1 |
| Fumarate | 3 | 2 | 2 | 2 | 2 | | 1 |
| Acetate | 4 | 21 | 5 | | | | 3 |
| L-malate | 2 | 2 | 1 | | 1 | | 7 |
| Formate | 34 | | | 40 | | | 5 |

*Fig S 10: The minimal subnetwork size for the uptake (up) and the secretion (down) of ten selected compounds in the seven-member bee gut microbiome and the number of alternative subnetworks.*

### Supplementary Tables

*Table S1: Number of metabolites (per model compartment, total and unique) in the $D_0$ networks of E. coli and S. cerevisiae.*

| Compartment in model | # of metabolites – *E. coli* | # of metabolites – *S. cerevisiae* |
|---|---|---|
| Cytoplasm | 83 | 59 |
| Periplasm | 15 | - |
| Mitochondrion | - | 36 |
| Mitochondrial Membrane | - | 1 |
| Lipid Particle | - | 2 |
| Peroxisome | - | 14 |
| Endoplasmic Reticulum | - | 1 |
| Total | 98 | 113 |
| Unique | 87 | 71 |

*Table S2: The cost of biosynthesis of biosynthesis of the common biomass building blocks for E. coli and S. cerevisiae. The metabolite abbreviations follow the BiGG[2] notation.*

| BBB | Cost of biosynthesis *E. coli* **:** Cost of biosynthesis *S. cerevisiae* |
|---|---|
| L-arginine | 4 nh4 + 4 nadph + akg + accoa + 3 atp + oaa + co2 **:** 4 nh4 + 2.5 nadph + akg + 6.5 atp + hco3 + oaa + 1.5 nadh |
| L-asparagine | h + 2 nh4 + nadph + atp + oaa **:** 2 nh4 + 0.5 nadph + 2.5 atp + oaa + 0.5 nadh + 0.5 h2o |
| L-aspartate | h + nh4 + nadph + oaa **:** nh4 + h + nadph + oaa |
| coenzyme A | 7 nh4 + 11 nadph + 0.83 pyr + accoa + 19.67 atp + 3 oaa + 2 3pg + 2 nad + prpp + so4 + 2 for + 1.17 pep **:** nh4 + nadph + akg |
| L-cysteine | 2 h + nh4 + 5 nadph + accoa + 2.33 atp + 3pg + nad + so4 + 0.33 pep **:** nh4 + h + 5 nadph + 2 atp + 3pg + accoa + nad + so4 |
| FAD | 10 nh4 + 4 nadph + 23.17 atp + 3 oaa + 2 co2 + 3 3pg + 4 nad + 2 prpp + 0.67 pep + 2 ru5p__D + 6.5 h2o **:** 13 nh4 + 5.5 nadph + 40.06 atp + 4 oaa + 1.5 nadh + 8 h2o + 3 co2 + 4 ru5p__D + 3 icit + 3 prpp + 0.06 fmn |
| L-glutamine | 2 nh4 + nadph + akg + 1.5 atp **:** 2 nh4 + 0.5 nadph + akg + 1.5 atp + 0.5 nadh |
| L-glutamate | h + nh4 + nadph + akg **:** nh4 + h + nadph + akg |
| L-glycine | nh4 + 3pg + nad + h2o **:** nh4 + h + nadph + icit |
| glycogen | atp + g6p **:** atp + h2o + g6p |
| L-histidine | 3 nh4 + 2 nadph + 4.33 atp + oaa + 2 nad + prpp + for + 0.33 pep + h2o **:** 4 nh4 + 1.5 nadph + 5.5 atp + oaa + 1.5 h2o + 3pg + 1.5 nad + 2 prpp + o2 |
| L-isoleucine | 5 h + nh4 + 5 nadph + pyr + 2 atp + oaa :nh4 + 2 nadph + pyr + akg |
| L-leucine | 2 h + nh4 + 2 nadph + 2 pyr + accoa + nad :nh4 + 2 h + 2 nadph + 2 pyr + accoa + nad |
| L-lysine | 4 h + 2 nh4 + 4 nadph + pyr + atp + oaa + succoa **:** 2 nh4 + h + 4 nadph + akg + atp + accoa + 2 nad |
| L-methionine | 4 h + 2 nh4 + 9 nadph + accoa + 3.33 atp + oaa + 2 3pg + nad + so4 + 0.33 pep + succoa **:** 2 nh4 + 5 h + 8.5 nadph + 3 atp + oaa + 3pg + accoa + 0.5 nad + so4 |
| NAD | 7.5 nh4 + 3 nadph + 12.08 atp + 3 oaa + 1.5 3pg + 0.5 nad + 2 prpp + 0.5 for + 0.33 pep + dhap + iasp **:** 8 nh4 + 4 nadph + 17 atp + 2 oaa + h2o + 2 3pg + icit + 4 prpp + 4 o2 + e4p + 2 pep |
| NADH | 7.5 nh4 + 3 nadph + 12.08 atp + 3 oaa + 1.5 3pg + 0.5 nad + 2 prpp + 0.5 for + 0.33 pep + dhap + iasp **:** 8 nh4 + 4 nadph + 17 atp + 2 oaa + h2o + 2 3pg + icit + 4 prpp + 4 o2 + e4p + 2 pep |

*Continued on the next page.*

| BBB | Cost of biosynthesis *E. coli* **:** Cost of biosynthesis *S. cerevisiae* |
|---|---|
| NADP | 7.5 nh4 + 3 nadph + 12.08 atp + 3 oaa + 1.5 3pg + 0.5 nad + 2 prpp + 0.5 for + 0.33 pep + dhap + iasp **:** 8 nh4 + 4 nadph + 17 atp + 2 oaa + h2o + 2 3pg + icit + 4 prpp + 4 o2 + e4p + 2 pep |
| NADPH | 7.5 nh4 + 3 nadph + 12.08 atp + 3 oaa + 1.5 3pg + 0.5 nad + 2 prpp + 0.5 for + 0.33 pep + dhap + iasp **:** 8 nh4 + 4 nadph + 17 atp + 2 oaa + h2o + 2 3pg + icit + 4 prpp + 4 o2 + e4p + 2 pep |
| L-phenylalanine | nh4 + 2 nadph + atp + 2 pep + e4p :nh4 + 2 h + 2 nadph + atp + e4p + 2 pep |
| L-proline | 2 h + nh4 + 3 nadph + akg + atp :nh4 + 3 h + 3 nadph + akg + atp |
| riboflavin | 4.5 nh4 + 2 nadph + 11.5833 atp + oaa + co2 + 1.5 3pg + 2.5 nad + prpp + 0.33 pep + 2 ru5p__D + 4.75 h2o **:** 13 nh4 + 5 nadph + 40 atp + 3 oaa + 11 h2o + 3 co2 + 4 ru5p__D + 3 icit + 3 prpp |
| L-serine | nh4 + nadph + 3pg + nad :nh4 + nadph + 3pg + nad |
| THF | 7.5 nh4 + 4 nadph + akg + 16.08 atp + oaa + co2 + 1.5 3pg + 3.5 nad + prpp + 2.33 pep + 7.25 h2o + e4p **:** 11 nh4 + 5.5 nadph + akg + 31.5 atp + 2 oaa + 0.5 nadh + 9.5 h2o + 2 co2 + 2 ru5p__D + 2 icit + 2 prpp + e4p + 2 pep |
| TDP | 5 nh4 + 9 nadph + 0.67 pyr + accoa + 13.83 atp + 2 3pg + 4 nad + 2 prpp + so4 + 2.33 pep + 2.5 h2o + e4p + g3p **:** 4 nh4 + 6.03 nadph + 13.47 atp + oaa + 2.52 3pg + 2 accoa + 0.83 nad + so4 + 2 icit + 2.52 prpp + 1.52 o2 + 0.51 r5p + 0.49 xu5p__D |
| L-threonine | 2 h + nh4 + 3 nadph + 2 atp + oaa :nh4 + 2 h + 2.5 nadph + 2 atp + oaa + 0.5 nadh |
| L-tryptophan | 2 nh4 + nadph + 2 atp + prpp + 2 pep + e4p **:** 2 nh4 + 1.5 nadph + 2.5 atp + 3pg + 0.5 nad + prpp + e4p + 2 pep |
| L-tyrosine | nh4 + 2 nadph + atp + nad + 2 pep + e4p :nh4 + h + nadph + atp + e4p + 2 pep |
| L-valine | 3 h + nh4 + 2 nadph + 2 pyr :nh4 + 3 h + 2 nadph + 2 pyr |

*Table S3: The in silico defined minimal media for the core bee gut microbiome.*

| *G. apicola* | *S. alvi* | *L. apis* | *L. kullabergensis* | *B. mellifer* | *B. mellis* | *B. asteroides* |
|---|---|---|---|---|---|---|
| D-Glucose | H2O | D-Glucose | D-Glucose | H2O | H2O | H2O |
| H2O | H+ | H2O | H2O | H+ | H+ | H+ |
| H+ | Chloride | H+ | H+ | L-Leucine | L-Leucine | Chloride |
| Chloride | Phosphate | L-Leucine | L-Leucine | Chloride | Chloride | Phosphate |
| Phosphate | Ammonium | Chloride | Chloride | Phosphate | Phosphate | Riboflavin |
| Ammonium | Fe3+ | Phosphate | Phosphate | Riboflavin | Riboflavin | Adenosine |
| Fe3+ | Potassium | Riboflavin | Ammonium | Ammonium | Ammonium | Ammonium |
| Potassium | Calcium | Aminoimidazole-riboside | L-Serine | L-Arginine | Guanine | L-Cysteine |
| Calcium | Citrate | L-Malate | L-Threonine | Cys-Gly | L-Serine | Potassium |
| Magnesium | Magnesium | L-Serine | L-Arginine | Fe3+ | L-Threonine | Calcium |
| Mn2+ | Mn2+ | L-Threonine | Cys-Gly | L-Aspartate | Cys-Gly | Magnesium |
| Co2+ | Co2+ | L-Arginine | Ornithine | L-Phenylalanine | Fe3+ | Mn2+ |
| Zinc | Zinc | Cys-Gly | L-Phenylalanine | Potassium | Ornithine | Co2+ |
| CMP | Cu2+ | Ornithine | Potassium | Beta-Alanine | L-Aspartate | Zinc |
| Cu2+ | O2 | L-Phenylalanine | Benzoate | Benzoate | Potassium | CMP |
| Nicotinate | Sulfate | Potassium | Calcium | Calcium | Beta-Alanine | Coenzyme A |
| Pyridoxal | Thiamin | Butyrate | Magnesium | Citrate | Calcium | Cu2+ |
| Sulfate | | Benzoate | Mn2+ | Magnesium | Magnesium | Fe2+ |
| Thiamin | | Calcium | Co2+ | Mn2+ | Mn2+ | D-Glucuronate |
| | | Magnesium | Zinc | Co2+ | Co2+ | Guanosine |
| | | Mn2+ | Cu2+ | Zinc | Zinc | Uaagmda |
| | | Co2+ | L-Glutamate | Cu2+ | Cu2+ | NMN |
| | | Zinc | DTMP | L-Glutamate | L-Glutamate | Shikimate |
| | | Cu2+ | Folate | Fe2+ | Fe2+ | Thiamin |
| | | L-Glutamate | Fumarate | Folate | Folate | Sulfate |

*Continued on the next page.*

| *G. apicola* | *S. alvi* | *L. apis* | *L. kullabergensis* | *B. mellifer* | *B. mellis* | *B. asteroides* |
|---|---|---|---|---|---|---|
| | | Deoxyadenosine | L-Histidine | L-Histidine | Cellobiose | Folate |
| | | Folate | L-Isoleucine | L-Methionine | L-Histidine | D-Mannose 1-phosphate |
| | | L-Histidine | L-Methionine | NMN | L-Isoleucine | |
| | | L-Methionine | NMN | Sulfate | L-Methionine | |
| | | Nicotinate | (R)-Pantothenate | L-Tryptophan | NMN | |
| | | (R)-Pantothenate | L-Tryptophan | L-Tyrosine | Sulfate | |
| | | L-Tryptophan | L-Tyrosine | L-Valine | L-Valine | |
| | | L-Tyrosine | D-Ribose | Thiamin | Shikimate | |
| | | L-Valine | Thiamin | Uracil | Thiamin | |
| | | Fe-enterobactin | Undecaprenyl-P | Xanthosine | Thymidine | |
| | | Sulfate | Uracil | | | |
| | | Thiamin | Xanthosine | | | |
| | | | Sulfate | | | |
| | | | Fe-enterobactin | | | |
| | | | Pyridoxal | | | |
| | | | L-Valine | | | |

*Table S4: The cost of biosynthesis of biosynthesis of the biomass building blocks for the core bee gut microbiome. The metabolite abbreviations follow the BiGG[2] notation. Empty entry signifies that the BBB can be produced from the core network.*

| BBB | Cost of biosynthesis *G. apicola* |
|---|---|
| 10fthf | 29 atp + 12 h2o + 8 nh4 + nad + 10 nadph + akg + co2 + 2 for + 2 pep + 3 oaa + r5p + e4p |
| ala_L | atp + nh4 + nadph + pyr |
| amet | 22.5 atp + 4.5 h2o + 6 nh4 + 10.33 nadph + co2 + accoa + 3pg + 2 for + 3 oaa + r5p + so4 + 0.33 nad |
| arg_L | 9 atp + 3 h2o + 4 nh4 + 4 nadph + akg + co2 + oaa |
| asn_L | 3.5 atp + 1.5 h2o + 2 nh4 + nadph + oaa |
| asp_L | atp + nh4 + nadph + oaa |
| coa | 14.75 atp + 6.75 h2o + nad + 5.5 nadph + 1.5 h_p + 1.5 h2o_p + pyr + 0.5 accoa + 3pg + for + oaa + 0.5 so4 + 1.5 cmp_p + nh4 |
| ctp | 6.667 atp + 3.67 h2o + h_p + h2o_p + cmp_p |
| cys__L | 4 atp + nh4 + h + nad + 5 nadph + accoa + 3pg + so4 |
| datp | 19 atp + 3 h2o + 5 nh4 + 0.66 coa + 5.33 nadph + 0.66 pyr + 2 for + 3 oaa + r5p + 0.33 co2 |
| dctp | 6.667 atp + 2.667 h2o + 0.66 coa + h_p + h2o_p + 0.66 pyr + cmp_p + 0.33 nadph |
| dgtp | 19.5 atp + 5.5 h2o + 5 nh4 + nad + 4.4 nadph + 0.4 co2 + 2 for + 2 oaa + r5p + 0.6 coa + 0.6 pyr |
| dttp | 5.667 atp + 0.66 h2o + 1.33 h + 3 nadph + h_p + h2o_p + for + cmp_p |
| fad | 39.5 atp + 13.5 h2o + 9 nh4 + nad + 10 nadph + 2 co2 + for + 2 ru5p__D + 5 oaa + 2 r5p |
| gln_L | 2 atp + 2 nh4 + nadph + akg |
| glu__L | atp + nh4 + nadph + akg |
| gly | 3 atp + h2o + nh4 + h + 3 nadph + oaa |
| gtp | 19.5 atp + 6.5 h2o + 5 nh4 + nad + 4 nadph + co2 + 2 for + 2 oaa + r5p |
| his_L | 9 atp + 4 h2o + 3 nh4 + 2 nad + 2 nadph + for + oaa + r5p |
| ile__L | 4 atp + nh4 + 3 h + 5 nadph + pyr + oaa |
| leu__L | atp + nh4 + h + nad + 2 nadph + 2 pyr + accoa |
| lys__L | 3 atp + 2 nh4 + 2 h + 4 nadph + pyr + succoa + oaa |
| met__L | 6 atp + nh4 + 6 h + 0.66 nadh + 8.33 nadph + accoa + for + oaa + so4 |
| mlthf | 29 atp + 11 h2o + 8 nh4 + nad + 11 nadph + akg + co2 + 2 for + 2 pep + 3 oaa + r5p + e4p |
| mql8 | 146.67 atp + 84.67 h2o + 25.33 nadh + 24.67 nadph + 14 pyr + 2 succoa + 2 for + 4 pep + 2 oaa + 2 r5p + 2 e4p + 16 g3p |
| nad | 24.5 atp + 8.5 h2o + 6 nh4 + 5 nadph + co2 + 2 for + 3 oaa + 2 r5p + nac_p |
| nadp | 4.9 atp + 1.7 h2o + 1.2 nh4 + nadph + 0.2 co2 + 0.4 for + 0.6 oaa + 0.4 r5p + 0.2 nac_p |
| phe__L | 2 atp + nh4 + h + 2 nadph + 2 pep + e4p |
| pro__L | 2 atp + nh4 + 2 h + 3 nadph + akg |

*Continued on the next page.*

| BBB | Cost of biosynthesis *G. apicola* |
|---|---|
| pydx5p | atp + pydx_p |
| ribflv | 19.5 atp + 8.5 h2o + 4 nh4 + nad + 5 nadph + co2 + 2 ru5p__D + 2 oaa + r5p |
| ser__L | atp + h2o + nh4 + nad + nadph + 3pg |
| thf | 28 atp + 12 h2o + 8 nh4 + nad + 10 nadph + akg + co2 + for + 2 pep + 3 oaa + r5p + e4p |
| thmpp | 4 atp + 2 h2o + thm |
| thr__L | 3 atp + h2o + nh4 + h + 3 nadph + oaa |
| tyr__L | 2 atp + nh4 + nad + 2 nadph + 2 pep + e4p |
| trp__L | 5 atp + 2 nh4 + nad + 2 nadph + 3pg + 2 pep + r5p + e4p |
| uaagmda | 100.33 atp + 52.33 h2o + 8 nh4 + 16.5 nadh + 20 nadph + 13.5 pyr + akg + succoa + 2 accoa + pep + 2.5 oaa + 2 f6p + 11 g3p |
| utp | 4.667 atp + 2.667 h2o + h_p + h2o_p + cmp_p |
| val__L | atp + nh4 + 2 h + 2 nadph + 2 pyr |

| BBB | Cost of biosynthesis *S. alvi* |
|---|---|
| 10fthf | 53.5 atp + 17.5 h2o + 15 nh4 + nad + 14 nadph + akg + 3 co2 + 3 prpp + 4 ru5p__D + 2 pep + 6 oaa + e4p |
| ala__L | atp + nh4 + nadph + pyr |
| amet | 53.5 atp + 4.5 h2o + 16 nh4 + 2 nad + 30 nadph + 3 co2 + 1.5 succoa + 3 3pg + 3 prpp + 4 ru5p__D + 7 oaa + 3 so4 + 1.5 accoa |
| arg__L | 8 atp + 2 h2o + 4 nh4 + 4 nadph + akg + co2 + oaa |
| asn__L | 2 atp + 2 nh4 + nadph + oaa |
| asp__L | atp + nh4 + nadph + oaa |
| coa | 46.75 atp + 11.25 h2o + 11.5 nh4 + nadh + 15.5 nadph + pyr + co2 + 0.5 accoa + 2.5 prpp + 4 ru5p D + 6.5 oaa + 0.5 so4 |
| ctp | 8 atp + 2 h2o + 3 nh4 + nadph + prpp + oaa + orot |
| cys__L | 2 atp + nh4 + 2 h + nad + 5 nadph + accoa + 3pg + so4 |
| datp | 49 atp + 13 h2o + 13 nh4 + nadh + 13 nadph + 3 co2 + 3 prpp + 4 ru5p__D + 7 oaa |
| dctp | 8 atp + h2o + 3 nh4 + 2 nadph + prpp + oaa + orot |
| dgtp | 49.5 atp + 14.5 h2o + 13 nh4 + 12 nadph + 3 co2 + 3 prpp + 4 ru5p__D + 6 oaa |
| dttp | 7 atp + 3 nh4 + nad + 4 nadph + 3pg + prpp + oaa + orot |
| fad | 50 atp + 14 h2o + 13 nh4 + nadh + 12 nadph + 3 co2 + 3 prpp + 4 ru5p__D + 7 oaa |
| gln__L | 2 atp + 2 nh4 + nadph + akg |
| glu__L | atp + nh4 + nadph + akg |
| gly | 3 atp + h2o + nh4 + h + nadh + 2 nadph + oaa |
| gtp | 49.5 atp + 15.5 h2o + 13 nh4 + 11 nadph + 3 co2 + 3 prpp + 4 ru5p__D + 6 oaa |

| | |
|---|---|
| his__L | 22.5 atp + 7.5 h2o + 7 nh4 + 2 nad + 6 nadph + co2 + 2 prpp + 2 ru5p__D + 3 oaa |

*Continued on the next page.*

| BBB | Cost of biosynthesis *S. alvi* |
|---|---|
| ile__L | 3 atp + nh4 + 3 h + nadh + 4 nadph + pyr + succoa + oaa |
| leu__L | atp + nh4 + h + nad + 2 nadph + 2 pyr + accoa |
| lys__L | 3 atp + 2 nh4 + 2 h + 4 nadph + pyr + succoa + oaa |
| met__L | 4 atp + 2 nh4 + 5 h + 8 nadph + 0.5 succoa + 3pg + oaa + so4 + 0.5 accoa |
| mlthf | 53.5 atp + 16.5 h2o + 15 nh4 + nad + 15 nadph + akg + 3 co2 + 3 prpp + 4 ru5p__D + 2 pep + 6 oaa + e4p |
| mql8 | 208 atp + 72 h2o + 4 nh4 + 44 nadh + 49 nadph + 28 pyr + 4 succoa + 4 3pg + 36 prpp + 8 pep + 4 e4p + 32 g3p |
| nad | 25.75 atp + 6.25 h2o + 7.5 nh4 + 0.5 nadh + 6.5 nadph + co2 + 2 prpp + 2 ru5p__D + 4 oaa + 0.5 dhap + 0.5 iasp |
| nadp | 3.96 atp + 0.96 h2o + 1.15 nh4 + 0.08 nadh + nadph + 0.15 co2 + 0.31 prpp + 0.31 ru5p__D + 0.61 oaa + 0.08 dhap + 0.08 iasp |
| phe__L | 2 atp + nh4 + h + 2 nadph + 2 pep + e4p |
| pro__L | 2 atp + nh4 + 2 h + 3 nadph + akg |
| pydx5p | atp + nh4 + 4 nad + nadph + pyr + e4p + g3p |
| ribflv | 16.5 atp + 5.5 h2o + 4 nh4 + 4 nadph + co2 + prpp + 2 ru5p__D + 2 oaa |
| ser__L | atp + h2o + nh4 + nad + nadph + 3pg |
| thf | 36 atp + 12 h2o + 11 nh4 + nad + 10 nadph + akg + 2 co2 + 2 prpp + 2 ru5p__D + 2 pep + 4 oaa + e4p |
| thmpp | 4 atp + h2o + thm |
| thr__L | 3 atp + h2o + nh4 + h + nadh + 2 nadph + oaa |
| trp__L | 3 atp + 2 nh4 + nad + 2 nadph + 3pg + prpp + 2 pep + e4p |
| tyr__L | 2 atp + nh4 + nad + 2 nadph + 2 pep + e4p |
| uaagmda | 83 atp + 23 h2o + 8 nh4 + 16.5 nadh + 20 nadph + 13.5 pyr + akg + succoa + 2 accoa + 11 prpp + pep + 2.5 oaa + 2 f6p + 11 g3p |
| utp | 6 atp + h2o + 2 nh4 + nadph + prpp + oaa + orot |
| val__L | atp + nh4 + 2 h + 2 nadph + 2 pyr |

| BBB | Cost of biosynthesis *L. apis* |
|---|---|
| 10fthf | 3.5 atp + 3.5 h2o + 0.5 h + 0.5 nadh + nadph + 0.5 pyr + 0.5 airs_p + 2.5 cgly + 0.5 g3p + fol + 0.5 tyr__L |
| ala__L | h2o + cgly |
| amet | 8.5 atp + 2 glu__L + 8 h2o + 0.5 nad + gtp + 3 ser__L + 2 oaa + 1.5 airs_p + 5 cgly + 0.5 g3p + 0.5 tyr__L + met__L_p |
| arg__L | |
| asn__L | 2 atp + glu__L + h2o + ser__L + oaa |
| asp__L | glu__L + ser__L + oaa |
| coa | 10.5 atp + 2 glu__L + 11 h2o + 0.5 nad + gtp + 3 ser__L + 2 oaa + 1.5 airs_p + 9 cgly + utp + 0.5 g3p + 0.5 tyr__L + pnto__R |

| | |
|---|---|
| ctp | 7.5 atp + 2 glu__L + 2.5 h2o + 2 ser__L + oaa + r5p + utp + orot |
| cys__L | h2o + cgly |

*Continued on the next page.*

| BBB | Cost of biosynthesis *L. apis* |
|---|---|
| datp | 3 atp + dad_2 |
| dctp | 8.5 atp + 2 glu__L + 2.5 h2o + nadph + 2 ser__L + oaa + r5p + utp + orot |
| dgtp | 9.5 atp + 8.5 h2o + 0.5 nad + nadph + 2 ser__L + oaa + 1.5 airs_p + 8.5 cgly + 0.5 g3p + 0.5 tyr__L |
| dttp | 8.5 atp + 3 glu__L + 2 nadph + 3 ser__L + oaa + r5p + orot + h2o + 0.5 utp |
| fad | 8.5 atp + 2 glu__L + 8 h2o + 0.5 nad + gtp + ribflv + 3 ser__L + 2 oaa + 1.5 airs_p + 7 cgly + 0.5 g3p + 0.5 tyr__L |
| gln__L | atp + 2 glu__L + ser__L + oaa |
| glu__L | |
| gly | h2o + cgly |
| gtp | 17 atp + 18.5 h2o + nad + 4 ser__L + 2 oaa + 3 airs_p + 16 cgly + g3p + tyr__L + 0.5 pep |
| his__L | |
| ile__L | 3 h + nadh + nadph + pyr + thr__L |
| leu__L | |
| lys__L | atp + 3 glu__L + 2 h2o + 3 ser__L + oaa + actp |
| met__L | atp + h2o + met__L_p |
| mlthf | |
| mql8 | 460 atp + 6 glu__L + 432 h2o + 455 h + 72 nad + 102 nadph + 16 gtp + 6 succoa + 11 fad + 96 ser__L + 240 crn + 72 but + 6 bz + 120 feenter_p + 6 met__L_p + 16 utp |
| nad | 23 atp + 24 h2o + 2 gtp + 2 ser__L + 4 oaa + 2 r5p + 3 airs_p + 18 cgly + g3p + tyr__L + 2 nac |
| nadp | 46 atp + 52 h2o + 5 gtp + 4 ser__L + 8 oaa + 4 r5p + 6 airs_p + 40 cgly + 2 g3p + 2 tyr__L + 4 nac |
| phe__L | |
| pro__L | h + nadh + pyr + orn |
| pydx5p | atp + 3 h2o + nad + r5p + 5 cgly + g3p |
| ribflv | |
| ser__L | |
| thf | 2 h + 2 nadph + fol |
| thmpp | 4.33 atp + 3 h2o + 0.33 gtp + thm_e + 0.33 utp |
| thr__L | |
| trp__L | |
| tyr__L | |

| | |
|---|---|
| uaagmda | 154 atp + 56.33 glu__L + 139 h2o + 85.67 h + 18 nad + 3.67 gtp + 45 ser__L + pep + oaa + 72 crn + 2 f6p + 18 but + 5.67 utp + 36 feenter_p |
| utp | 14 atp + 6 glu__L + 4 h2o + 4 ser__L + 2 oaa + 2 r5p + 2 orot + pep |
| val__L | |

*Continued on the next page.*

| BBB | Cost of biosynthesis *L. kullabergensis* |
|---|---|
| 10fthf | 4 atp + 1.67 h2o + 0.67 h + 2.33 nadph + 0.67 ru5p__D + 1.17 oaa + 0.67 rib__D + fol + 2.33 trp__L + 0.33 xtsn + 1.17 pyr |
| ala__L | h2o + cgly |
| amet | 8 atp + 5 h2o + nadph + oaa + met__L + trp__L + xtsn |
| arg__L | |
| asn__L | 2 atp + 2 nh4 + nadh + oaa |
| asp__L | nh4 + h + nadh + oaa |
| coa | 12 atp + 13457.6 h2o + nadph + oaa + 13452.6 cgly + trp__L + xtsn + pnto__R |
| ctp | 6 atp + h2o + nh4 + rib__D + ura |
| cys__L | h2o + cgly |
| datp | 8 atp + 2 h2o + 2 nadph + 2 oaa + 2 trp__L + xtsn |
| dctp | 6 atp + nh4 + nadph + 0.5 pyr + rib__D + ura + trp__L + 0.5 oaa |
| dgtp | 7 atp + 2 h2o + nh4 + nadph + 0.5 pyr + trp__L + xtsn + 0.5 oaa |
| dttp | 2 atp + h + dtmp |
| fad | 18 atp + 9 h2o + 2 nadph + 2 ru5p__D + 2 oaa + 2 rib__D + 2 trp__L + 2 xtsn |
| gln__L | atp + glu L + nh4 |
| glu__L | |
| gly | h2o + cgly |
| gtp | 7 atp + 3 h2o + nh4 + xtsn |
| his_L | |
| ile_L | |
| leu__L | |
| lys__L | atp + 4 h + 2 nadh + pyr + oaa + actp + 2 trp__L |
| met__L | |
| mlthf | 4 h + 2 nadph + 0.67 pyr + ser__L + fol + 2 trp__L + 1.33 oaa |
| mql8 | 31 atp + 62.5 h2o + 35.5 nad + 16 nadph + 11.5 pyr + succoa + ser__L + 30 glyc3p + bz + 12.5 val__L + fum |
| nad | 8 atp + 4 h2o + nadph + oaa + nmn + trp__L + xtsn |
| nadp | 8 atp + 4 h2o + nadph + oaa + nmn + trp__L + xtsn |

| | |
|---|---|
| phe__L | |
| pro__L | orn |
| pydx5p | atp + pydx_e |
| ribflv | 9 atp + 5 h2o + nadph + 0.5 pyr + 2 ru5p__D + 2 rib__D + trp__L + xtsn + 0.5 oaa |

*Continued on the next page.*

| BBB | Cost of biosynthesis *L. kullabergensis* |
|---|---|
| ser_L | |
| thf | 4 h + 2 nadph + pyr + fol + 2 trp_L + oaa |
| thmpp | 4 atp + 2 h2o + thm |
| thr_L | |
| trp_L | |
| tyr__L | |
| uaagmda | 11 atp + glu__L + 92.36 nh4 + 90.36 nadh + 90.36 pyr + pep + udcpp + oaa + actp + 2 f6p + val__L + 81.36 h |
| utp | 5 atp + h2o + rib__D + ura |
| val__L | |

| BBB | Cost of biosynthesis *B. mellifer* |
|---|---|
| 10fthf | 2 h2o + h + nad + nadph + cgly + fol |
| ala_L | h2o + cgly |
| amet | 8.1 atp + 5.1 h2o + nadph + asp__L + met__L + 1.1 xtsn |
| arg_L | |
| asn__L | 2 atp + h2o + nh4 + asp__L |
| asp__L | |
| coa | 19.30 atp + 8.30 h2o + 3 nadph + pyr + asp__L + actp + ala_B + cgly + ura + val__L + 2.3 xtsn + 0.5 nadh |
| ctp | 6 atp + 2 h2o + nh4 + ura + xtsn |
| cys__L | h2o + cgly |
| datp | 8.2 atp + 2.2 h2o + 2 nadph + asp__L + 1.2 xtsn |
| dctp | 7.1 atp + 2.1 h2o + nh4 + nadph + 1.1 xtsn |
| dgtp | 7.1 atp + 2.1 h2o + nh4 + nadph + 1.1 xtsn |
| dttp | 5.1 atp + 2.1 h2o + 1.67 nadph + cgly + ura + 1.1 xtsn + 0.67 nad |
| fad | 9.1 atp + 4.1 h2o + nadph + asp__L + ribflv + 1.1 xtsn |
| gln__L | atp + glu__L + nh4 |
| glu__L | |

| | |
|---|---|
| gly | h2o + cgly |
| gtp | 7 atp + 3 h2o + nh4 + xtsn |
| his__L | |
| ile_L | 2.6 atp + glu__L + 5.8 h + 4.5 nadh + 2.5 nadph + 2 pyr + asp__L + 2 actp + 0.6 xtsn |
| leu_L | |

*Continued on the next page.*

| BBB | Cost of biosynthesis *B. mellifer* |
|---|---|
| lys_L | 1.3 atp + glu__L + 0.3 h2o + 2.9 h + 1.5 nadh + 2 nadph + pyr + asp__L + 2 actp + 0.3 xtsn |
| met_L | |
| mlthf | 4 h2o + 2 h + 2 nad + 2 nadph + 3 cgly + fol |
| mql8 | 51.14 atp + glu__L + 29.59 h2o + 2.21 nad + 19.88 nadph + 0.07 h2o_p + pyr + 24 actp + 1.14 cgly + bz + 2.74 ura + 6.80 xtsn + 1.59 nh4 |
| nad | 8.1 atp + 4.1 h2o + nadph + asp__L + nmn + 1.1 xtsn |
| nadp | 8.1 atp + 4.1 h2o + nadph + asp__L + nmn + 1.1 xtsn |
| phe_L | |
| pro__L | |
| pydx5p | 2.6 atp + nh4 + g3p + 1.60 xtsn |
| ribflv | |
| ser__L | 3 h2o + nad + nadph + 2 cgly |
| thf | 0.2 atp + 0.2 h2o + 1.6 h + 2 nadph + fol + 0.2 xtsn |
| thmpp | 4 atp + 2 h2o + thm |
| thr_L | 2.2 atp + 1.2 h2o + 0.6 h + 1.5 nadph + asp__L + 0.2 xtsn + 0.5 nadh |
| trp_L | |
| tyr__L | |
| uaagmda | 53 atp + 4 glu__L + 35 h2o + nh4 + 0.5 nad + 25.5 nadph + 4 pyr + asp__L + pep + 36 actp + 2 f6p + 2 cgly + 8 xtsn |
| utp | 5 atp + 2 h2o + ura + xtsn |
| val__L | |

| BBB | Cost of biosynthesis *B. mellis* |
|---|---|
| 10fthf | h2o + nad + nadp + gly + thf |
| ala__L | h2o + cgly |
| amet | 5 atp + 3 h2o + nadph + asp__L + r5p + met__L + gua |
| arg_L | 5 atp + 3 h2o + nh4 + co2 + asp__L + orn |
| asn__L | 2 atp + h2o + nh4 + asp__L |
| asp__L | |

| | |
|---|---|
| coa | 13 atp + 6.5 h2o + 1.5 nad + 3 nadph + 2 h_p + asp__L + pep + 2 r5p + cgly + ala_B_p + utp + val__L + gua + skm_p |
| ctp | 11 atp + 4 glu__L + 0.5 nad + 4 nadph + r5p + utp + thymd |
| cys__L | h2o + cgly |
| datp | 5 atp + 2 nadph + asp__L + r5p + gua |
| dctp | 11 atp + 4 glu_L + 0.5 nad + 5 nadph + r5p + utp + thymd |

*Continued on the next page.*

| BBB | Cost of biosynthesis *B. mellis* |
|---|---|
| dgtp | 4 atp + nadph + r5p + gua |
| dttp | 3 atp + thymd |
| fad | 6 atp + 2 h2o + nadph + asp__L + ribflv + r5p + gua |
| gln_L | atp + glu__L + nh4 |
| glu_L | |
| gly | |
| gtp | 4 atp + h2o + r5p + gua |
| his_L | |
| ile_L | |
| leu__L | |
| lys__L | atp + 3 h + 2.5 nadph + pyr + asp_L + actp + orn + 0.5 nadh |
| met__L | |
| mlthf | nad + gly + thf |
| mql8 | 26 atp + glu__L + 15.5 h2o + 24.5 nad + 16 nadph + 2 h_p + 11 acald + 2 pep + r5p + 2dmmql8 + 4 utp + thymd + 2 skm_p + 11 pyr |
| nad | 5 atp + 2 h2o + nadph + asp__L + r5p + nmn + gua |
| nadp | 5 atp + 2 h2o + nadph + asp__L + r5p + nmn + gua |
| phe_L | atp + 0.5 nadph + h_p + pep + orn + skm_p + 0.5 nadh |
| pro_L | atp + h_p + pep + orn + skm_p + 0.5 nadph |
| pydx5p | atp + nh4 + r5p + g3p |
| ribflv | |
| ser_L | |
| thf | |
| thmpp | 3.5 atp + 2 h2o + 0.5 utp + thm |
| thr__L | |
| trp_L | 4 atp + nh4 + h_p + ser__L + pep + r5p + skm_p |

| | |
|---|---|
| tyr__L | atp + 0.5 nad + 0.5 nadph + h_p + pep + orn + skm_p |
| uaagmda | 36.5 atp + glu__L + 26 h2o + 2 nh4 + 34.5 nad + 25.5 nadph + 18.5 pyr + asp__L + 17.5 acald + pep + actp + orn + 2 f6p + 3 cgly + 7.5 utp |
| utp | 40 atp + 16 glu__L + 3 h + 2 nad + 16 nadph + 4 r5p + + 4 thymd + pep |
| val__L | |

| BBB | BBB Cost of biosynthesis *B. asteroides* |
|---|---|
| 10fthf | atp + 2 h + 2 nadph + for + fol |

*Continued on the next page.*

| BBB | BBB Cost of biosynthesis *B. asteroides* |
|---|---|
| ala__L | nh4 + h + nadph + pyr |
| amet | 6 atp + 0.33 h2o + 2.67 h + 1.17 nadh + 5.5 nadph + adn + succoa + for + 2.67 oaa + cys__L_p + 1.67 nh4 |
| arg__L | 7.5 atp + 4 nh4 + 4 nadph + akg + co2 + oaa + 1.5 h2o |
| asn__L | 3.5 atp + 1.5 h2o + 2 nh4 + nadph + oaa |
| asp__L | nh4 + h + nadph + oaa |
| coa | |
| ctp | 2 atp + h + cmp |
| cys__L | atp + h2o + cys__L_p |
| datp | 3 atp + h + nadph + adn |
| dctp | 2 atp + 2 h + nadph + cmp |
| dgtp | 7 atp + 4 h2o + nad + nadph + adn |
| dttp | 4 atp + 5.5 h + 3 nadph + for + 1.5 cmp |
| fad | 4 atp + h2o + adn + ribflv |
| gln__L | 1.5 atp + 2 nh4 + nadph + akg |
| glu__L | nh4 + h + nadph + akg |
| gly | 2 atp + 3 nh4 + 4 h + 0.5 nadh + 4.5 nadph + 3 oaa |
| gtp | 7 atp + 5 h2o + nad + adn |
| his__L | 8 atp + 3 h2o + 3 nh4 + 2 nad + 2 nadph + for + oaa + r5p |
| ile__L | 2 atp + 8.5 nh4 + 12.5 h + 12 nadph + pyr + 8.5 oaa + 0.5 nadh |
| leu__L | nh4 + 2 h + nad + 2 nadph + 2 pyr + accoa |
| lys__L | atp + 8 nh4 + 10 h + 10 nadph + pyr + succoa + 7 oaa |
| met__L | 3 atp + 4.67 h + 1.16667 nadh + 5.5 nadph + succoa + for + 2.67 oaa + cys__L_p + 1.67 nh4 |
| mlthf | atp + 3 h + 3 nadph + for + fol |
| mql8 | 1536 atp + 192 h2o + 480 h + 608 nadh + 448 nadph + 335 pyr + 48 akg + 48 for + 48 pep + 48 r5p + 384 g3p + 48 skm + lac__L |
| nad | 3 atp + h2o + adn + nmn |

| | |
|---|---|
| nadp | 7 atp + 2 h2o + 2 adn + 2 nmn |
| phe__L | atp + nh4 + h + nadph + pep + skm |
| pro__L | atp + nh4 + 3 h + 2 nadph + akg + nadh |
| pydx5p | atp + nh4 + r5p + g3p |
| ribflv | |
| ser__L | 3 atp + 4.5 nh4 + 6.5 h + 0.5 nadh + 7 nadph + for + 4.5 oaa |
| thf | 2 h + 2 nadph + fol |
| thmpp | 4 atp + 2 h2o + thm |
| thr__L | 2 atp + 3 nh4 + 4 h + 4.5 nadph + 3 oaa + 0.5 nadh |
| trp__L | 11 atp + 9 nh4 + 0.5 nadh + 10.5 nadph + for + pep + 8 oaa + r5p + skm + 2 h |
| tyr__L | atp + nh4 + nad + nadph + pep + skm |
| uaagmda | |
| utp | 2 atp + h2o + 2 h + cmp |
| val__L | nh4 + 3 h + 2 nadph + 2 pyr |
